# Population genomics of the mostly thelytokous *Diplolepis rosae* (Linnaeus, 1758) (Hymenoptera: Cynipidae) reveals population-specific selection for sex

**DOI:** 10.1101/2023.01.26.525659

**Authors:** Ksenia Mozhaitseva, Zoé Tourrain, Antoine Branca

**Affiliations:** Laboratoire Evolution, Génomes, Comportement, Ecologie, l’Institut Diversité, Ecologie et Evolution du Vivant, Université Paris-Saclay, Gif-sur-Yvette, France

**Keywords:** *Diplolepis rosae*, thelytoky, recombination, runs of homozygosity, selection, population genomics

## Abstract

In Hymenoptera, arrhenotokous parthenogenesis (arrhenotoky) is a common reproductive mode. Thelytokous parthenogenesis (thelytoky), when virgin females produce only females, is less common and is found in several taxa. In our study, we assessed the efficacy of recombination and the effect of thelytoky on the genome structure of *Diplolepis rosae*, a gall wasp producing bedeguars in dog roses. We assembled a high-quality reference genome using Oxford Nanopore long-read technology and sequenced 17 samples collected in France with high-coverage Illumina reads. We found two *D*. *rosae* peripatric lineages that differed in the level of recombination and homozygosity. The first *D*. *rosae* lineage showed a recombination rate that was 13.2 times higher and a per-individual heterozygosity that was 1.6 times higher. We inferred that genes under negative selection were enriched in functions related to male traits (‘sperm competition’, ‘insemination’, and ‘copulation’ gene ontology terms) in the more recombining lineage, while in the less recombining form, the same lineage genes showed traces pointing towards balancing or relaxed selection. Thus, although *D*. *rosae* reproduces mainly by thelytoky, selection may act to maintain sexual reproduction.

**Significance:** Many organisms can alternate between sexual and asexual reproduction in different ways. Sexual reproduction is essential to creating genetic diversity for adaptation to changing environments, whereas asexual reproduction is important in the short term and in stable environments. Using genomic data, we demonstrated the existence of two lineages in the rose bedeguar wasp *Diplolepis rosae* previously shown to reproduce mainly by thelytokous parthenogenesis, giving almost only females. One of the lineages showed higher recombination, higher heterozygosity, and genes involved in male traits under negative selection. This could be linked to the expected advantages of maintaining sexual reproduction in natural populations.

## Introduction

It is commonly asserted that sexual reproduction, i.e. recombination, is advantageous because it provides the opportunity for organisms to generate genetic diversity, enabling them to overcome environmental perturbations. However, recombination can also break linked advantageous alleles and broaden the variance around a fitness optimum, leading to the counterselection of sexual lineages (Crow and Kimura 1965). Therefore, the coexistence of sexually and asexually reproducing forms, which exhibit variability in the frequency of recombination and heterozygosity, is related to the cost of sex from both short- and long-term perspectives (Williams 1966; Maynard-Smith 1978; Hamilton 1980). Species that possess both sexual and asexual lineages are especially interesting for studying the conditions that determine their selection for recombination.

In Hymenoptera, some species reproduce both sexually and asexually. Arrhenotokous parthenogenesis (arrhenotoky) is the ancestral sexual reproductive mode (Heimpel and De Boer 2008; Rabeling and Kronauer 2013). Diploid females develop from fertilised eggs, while haploid males develop from unfertilised eggs. Another less frequent mode of reproduction is thelytokous parthenogenesis (thelytoky), which relates to asexuality, in which virgin females produce only females (Heimpel and De Boer 2008). Thelytoky can be encoded in the genomes of hymenopterans (Wenseleers and Billen 2000; Belshaw and Quicke 2003; Engelstädter et al. 2011; Foray et al. 2013; Capdevielle-Dulac et al. 2022) or induced by endosymbionts (Stouthamer et al. 1990; Stouthamer and Kazmer 1994). Genetically based thelytoky exists in the form of automixis, i.e. gamete fusion or gamete duplication after meiosis, and in the form of apomixis, i.e. mitotic division of ootids with no meiosis (Heimpel and De Boer 2008; Schön et al. 2009; Queffelec et al. 2021). Notably, automictic Hymenopteran females can still produce rare males (Stenberg and Saura 2009). The presence of males in thelytokous populations can also be explained by a failure of the mechanism inducing thelytoky or by occasional gene flow between the arrhenotokous and thelytokous populations (Stouthamer and Kazmer 1994; Engelstädter et al. 2011). Thelytoky leads to a decrease in recombination and individual genetic diversity compared to arrhenotoky; therefore, each mode of reproduction will leave a contrasting pattern of genetic diversity across the genome (Tvedte et al. 2019).

Endosymbiont-induced thelytoky is caused by endocytoplasmic bacteria, such as *Rickettsia*, *Cardinium*, and *Wolbachia* (Adachi-Hagimori et al. 2011; Werren et al. 2008; Giorgini et al. 2009). *Wolbachia* is the most widespread intracellular parasite infecting 25 to 70% of all insect species (Kozek and Rao 2007). *Wolbachia*-induced thelytoky has been extensively studied in *Trichogramma* spp. (Stouthamer et al. 1990; Stouthamer and Kazmer 1994), where thelytokous females exposed to antibiotic treatment or high temperatures produced males and arrhenotokous females (Stouthamer et al. 1990). Thus far, *Wolbachia*-mediated thelytoky has only been shown to be induced via gamete duplication, i.e. the failure of chromosome segregation in unfertilised eggs during anaphase I and subsequent duplication of terminal meiotic products. This mechanism leads to completely homozygous females (Stouthamer and Kazmer 1994). Hence, endosymbiont-induced thelytoky may result in the rapid spread of several locally adapted parthenogenetic lineages, each originating from a single female, which might lead to speciation (Werren 1998; Schilder et al. 1999; Bordenstein 2003; Adachi-Hagimori et al. 2011).

*Diplolepis rosae* (Hymenoptera: Cynipidae) is a cynipid wasp that parasitises wild dog roses (*Rosa* spp. section *caninae*) and causes specific plant outgrowths called rose bedeguar galls (Shorthouse and Floate 2010; Giron et al. 2016). One cytological study of meiosis in *D*. *rosae* showed that it reproduces by thelytokous parthenogenesis via gamete duplication; after anaphase II, one of the haploid ootids enters mitotic division. Thereafter, the fusion of the two daughter products results in the restoration of diploidy, thereby leading to completely homozygous females (Stille and Dävring 1980). Indeed, *D*. *rosae* individuals collected from several European locations consisted of almost only females, with the proportion of males varying from 1 to 4% (Nordlander 1973). A further study based on the electrophoresis of 27 isozymes showed that *D*. *rosae* females sampled in Sweden, Germany, and Greece were completely homozygous (Stille 1985). Additionally, females of *D*. *rosae* have been shown to be infected with *Wolbachia* (Plantard et al. 1999; Schilthuizen and Stouthamer 1998; Van Meer et al. 1995). For instance, in France, the percentage of *Wolbachia*-infected females varied from 71 to 100%, depending on the location (Plantard et al. 1999). Finally, *Wolbachia*-induced thelytoky was indirectly confirmed by examining three microsatellite loci in a closely related species, *D*. *spinosissimae* (Zhang et al. 2020). In *Wolbachia*-infected populations, males comprised about 1.3% of individuals, and no heterozygous females were found. Conversely, in *Wolbachia*-free populations, males represented 21–29% of individuals, and 78–96% of females were heterozygous (Plantard et al. 1998).

In summary, the previously published *D*. *rosae* studies demonstrated that populations are strongly female-biased. Based on several genetic markers, females were mostly homozygous. The prevalence of females and homozygosity associated with *Wolbachia* infection were supposed to induce thelytokous parthenogenesis. Therefore, we expected that *Wolbachia* infection in *D*. *rosae* might lead to gamete duplication thelytoky and a fine-scale population structure coupled with high homozygosity across the genome. However, any selection for recombination and genetic diversity would act to maintain sexual reproduction and leave different signatures on the genome.

Thus, the objective of our study was to assess the effect of thelytoky on the genome of *D*. *rosae*. First, using a high-quality reference genome of *D*. *rosae*, we investigated whether thelytoky led to a fine-scale population structure of *D*. *rosae* in France. Second, we analysed the patterns of diversity, homozygosity, and recombination to show the consequences of thelytoky on the genome of *D*. *rosae*. Third, we assessed whether selection favoured recombination over thelytoky by searching for regions that showed specific patterns of differentiation, recombination, and homozygosity.

## Results

### Genome structure

The total length of the *D*. *rosae* reference genome assembly was estimated at 760.7 Mbp, with a total number of sequences of 757, an N50 of 7,663,408 bp, the largest scaffold of 33,454,033 bp, and an L50 of 25 (**Table S1**). Repetitive sequences represented 69.28% of the genome assembly: 48.50% unclassified repeats, 11.40% retroelements, 8.66% DNA transposons, and 0.72% other repeats (rolling circles, small RNA, satellites, simple repeats, and small complexity repeats) (**Table S2**). Using the BUSCO hymenoptera_odb10 dataset (Manni et al. 2021), we found a genome completeness of 91.8% of complete and single-copy genes, 0.5% of complete and duplicated genes, 1.6% of fragmented genes, and 6.1% of missing genes. The total number of genes predicted by BRAKER2 (Hoff et al. 2019) *in silico* was 20,301, of which 14,559 were partially or fully annotated. The assembled genome of *D*. *rosae* is available at DDBJ/ENA/GenBank under accession JAPYXD000000000.

### Population structure

The most likely population structure for *D*. *rosae* in France was two lineages, population 1 and population 2. This was based on the number of lineages examined (K), which was equal to 2 (Raj et al. 2014) (**Fig. 1**). One individual, *D*. *rosae*-652, showed admixed ancestry, with 86.2% of its genome assigned to population 2 and 13.8% to population 1. The negative value of F3-admixture statistics (Patterson et al. 2012; Peter 2016) (F3 = −0.0797 ± 0.0150, z = −5.29, p < 0.0001) suggests that this individual is admixed between lineages related to population 1 and population 2. The additional genotyping confirmed two lineages distributed in France (**Fig. 2, Fig. S1**).

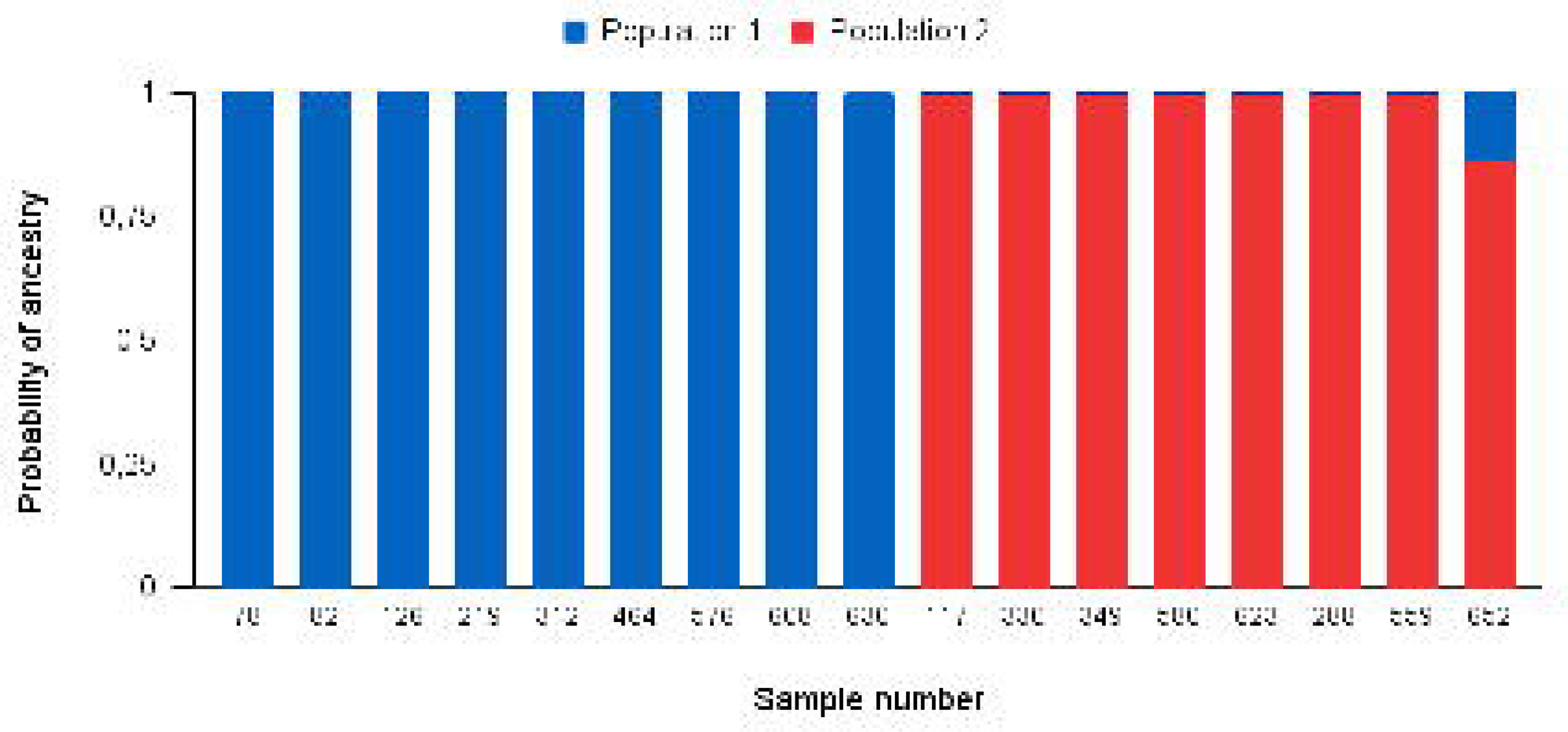

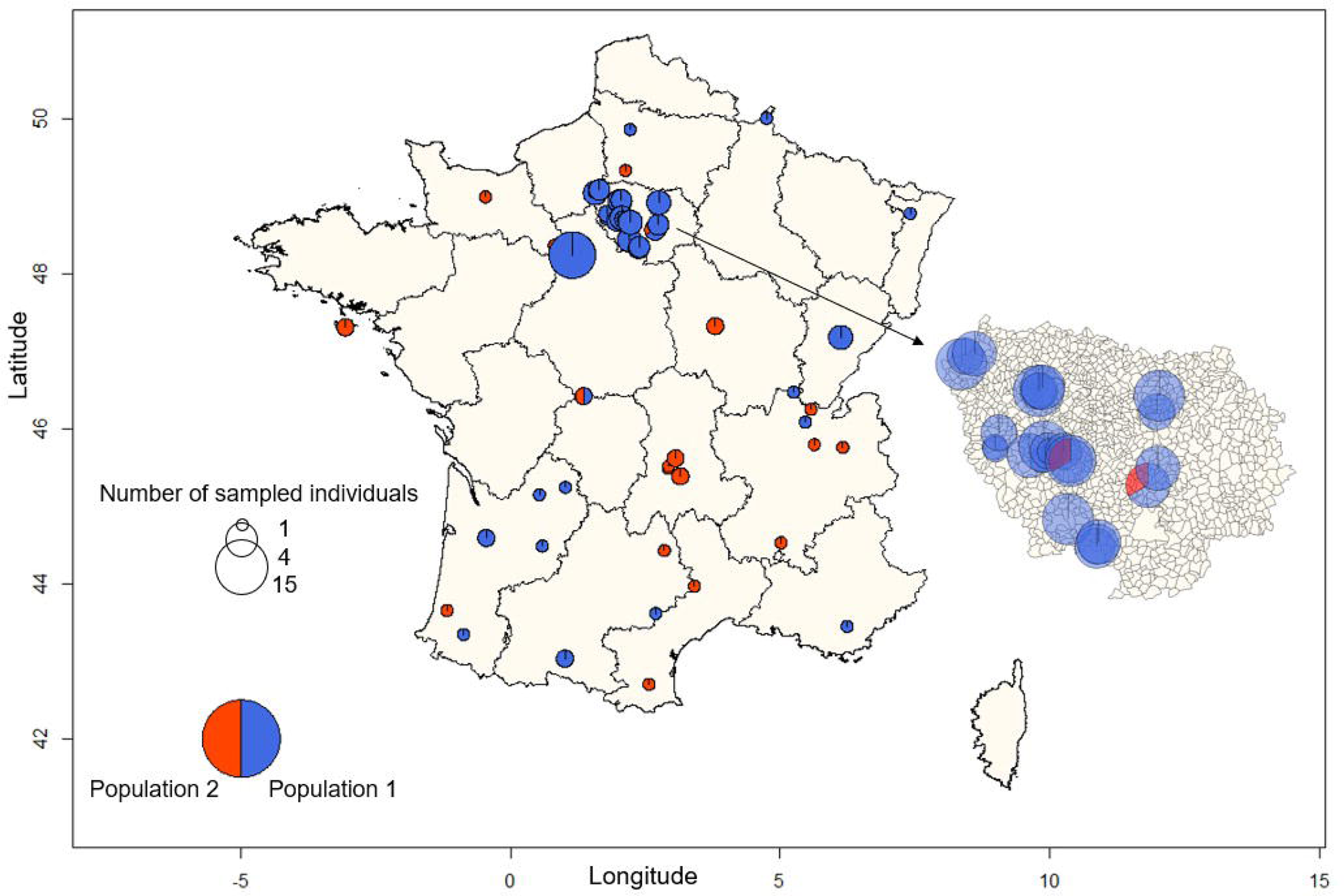

### Demographic scenario

According to the inference method (*dadi*) based on the joint site frequency spectrum of genetic variants (Gutenkunst et al. 2009), the most probable demographic scenario for *D*. *rosae* was a bottleneck of an ancestral population, followed by exponential growth, then a split into two populations with no gene flow between them (lowest AIC = 74419.2, **Table S3**, **Fig S2**). The bottleneck time and split time were estimated at 0.22*2Neff generations ago and 0.20*2Neff generations ago, respectively, where Neff is the effective population size (diploid individuals). The likelihood ratio test showed no significant difference between the models with and without the migration parameter (Dadj = 0.37, p = 0.28). The migration rate (m) was estimated as 0.81/(2*Neff) migrants per generation. Additional examination by approximate Bayesian computation (ABC) (Csilléry et al. 2012) confirmed the demographic model inferred by *dadi* to provide a good fit to the data (1 million simulations, Goodness of Fit, dist. = 3365.0, p = 0.21). The prediction errors for all model parameters (effective population size, bottleneck time, and split time) were close to 1 (**Table S4**). Posteriors had the same or broader distributions as the priors, except for effective population sizes (**Fig. S3**). The population sizes for both *D*. *rosae* lineages were estimated to be 500 diploid individuals.

### Population genomic statistics

The fixation index Fst and the absolute divergence Dxy values between the two populations were 0.81 ± 0.25 and 0.0033 ± 0.0024, respectively. Population 1 was characterised by a population-scaled recombination rate (ρ) that was 13.2 times higher (U = 511,350, z = 61.8, p = 0.0001) and a heterozygosity (H) that was 1.6 times higher (U = 9, z = 2.55, p = 0.0108) than population 2 (**Fig. 3**). In population 1, the median Tajima’s D value did not differ from zero (one-sample Wilcoxon test, comparison with the median u = 0: W = 9.5e+08, p = 0.22) (**Fig. 3**). In population 2, the distribution of Tajima’s D showed a slight excess of negative values (Tajima’s D median = −0.84, comparison with the median u = 0: W = 5.5e+08, p < 0.0001). There was no correlation between nucleotide diversity and recombination rate in population 1 (Pearson’s correlation r = 0.0626, t = 0.259, p = 0.799) or population 2 (r = 0.0287, t = 0.119, p = 0.907). There was no correlation between the protein-coding sequence density and nucleotide diversity in population 1 (r = −0.0142, t = −0.218, p = 0.827) or population 2 (r = −0.0237, t = −0.365, p = 0.715) (**Fig. S4**).

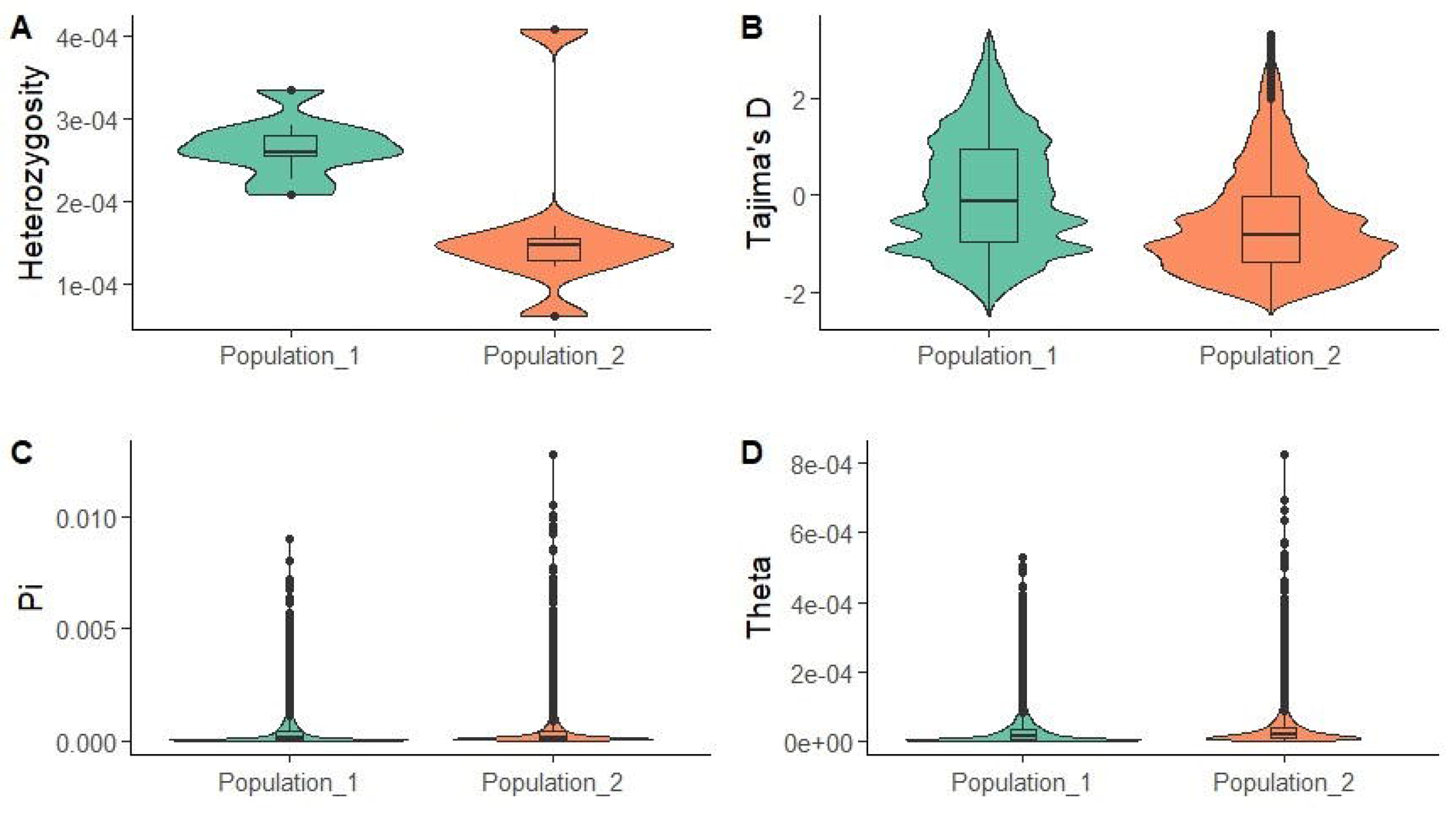

### Runs of homozygosity

Population 1 showed more frequent short (0.01–0.1 Mbp) runs of homozygosity (ROHs) than population 2 (Mann–Whitney U test: U = 9, z = 2.55, p = 0.0108) (**Fig. S5**). In contrast, population 2 showed more widespread and larger ROHs (0.1– 0.5 Mbp ROHs: U = 9, z = 2.454, p = 0.0141; 0.5–1.0 Mbp ROHs: U = 9, z = 2.550, p = 0.0108; >1 Mbp ROHs: U = 9, z = 2.550, p = 0.0108). In population 1, the ROHs covered 74 to 82% of the genome assembly length, whereas in population 2, the coverage varied from 83 to 91%. In the admixed individual, *D*. *rosae*-652 (**Fig. 1**), the ROHs covered 52% of the genome.

### Detection of homozygous genomic regions with low differentiation

In several regions of the *D*. *rosae* genome, longer ROHs overlapped with a decrease in Fst and a change in genetic diversity (π) (**Fig. 4**). Therefore, we developed a composite score (CS) that summarises Fst and π. It enabled us to detect the regions showing a drop in Fst with a simultaneous increase (positive CS) or decrease (negative CS) in π. We detected a positive CS (**Table 1**) associated with an increase in recombination rate in scaffold 204 (26.0 and 28.0 Mbp regions) (**Fig. 5**), scaffold 325 (5.1–7.5 Mbp) (**Fig. 6**), and scaffold 414 (5.5–9.0 Mbp) (**Fig. 6**) in population 1: some alleles segregated in both lineages but recombination broke the linkage between alleles in population 1. In scaffold 313 (6.1–6.2 Mbp) (**Fig. 5**), scaffold 325 (2.5–3.0 Mbp) (**Fig. 6**), and scaffold 762 (12.5–20.0 Mbp) (**Fig. 7**), the positive CS values (**Table 1**) in both lineages showed an increase in diversity π but a decrease in recombination rate ρ, indicating several haplotypes segregating in both populations with complete linkage. Furthermore, in population 1, scaffold 325 (2.5–3.0 Mbp Mbp) (**Fig. 6**) and scaffold 414 (5.8–6.0 Mbp) (**Fig. 6**) demonstrated a higher π*_N_*/π_S_ ratio, indicating an increase in non-synonymous site diversity relative to synonymous site diversity. In scaffold 204 (30.0 Mbp) (**Fig. 5**), scaffold 313 (4.1 Mbp) (**Fig. 5**), and scaffold 414 (3.0 Mbp) (**Fig. 6**), we detected regions with negative CS outliers (**Table 1**) associated with a lower recombination rate ρ: some linked alleles segregated in both *D*. *rosae* lineages. In scaffold 204 (30.1–33.0 Mbp) (**Fig. 5**), scaffold 313 (0.0–4.0 Mbp and 4.2–5.9 Mbp), and scaffold 523 (0.0–2.0 Mbp) (**Fig. 7**), we observed the opposite trend with the CS values (**Table 1**): positive outliers in population 1 but negative outliers in population 2. Gene set enrichment analysis (GSEA) of the genome regions showing outlier composite score values revealed several significantly enriched Gene Ontology (GO) terms (**Table S5**). In ‘Biological Process’ GO, the term ‘commissural neuron axon guidance’ corresponded to a negative score outlier in population 2 and a positive score outlier in population 1 (**Table S5**, **Fig. S6**). The terms ‘sperm competition’, ‘insemination’, and ‘copulation’ corresponded to a negative score outlier in population 1 (**Table S5**, **Fig. S7**) and a positive score outlier in population 2. In ‘Molecular Function’ GO, the terms ‘metalloendopeptidase activity’ and ‘metallopeptidase activity’ corresponded to negative outliers in population 1 (**Table S5**, **Fig. S8**) and positive outliers in population 2. The term ‘commissural neuron axon guidance’ annotated a group of genes found in scaffold 313 (1.7 – 2.7 Mbp region) (**Fig. 5**). The terms ‘sperm competition’, ‘insemination’, ‘copulation’, ‘metalloendopeptidase activity’, and ‘metallopeptidase activity’ matched genes found in scaffold 204 (28.7–28.9 Mbp region) (**Fig. 5**).

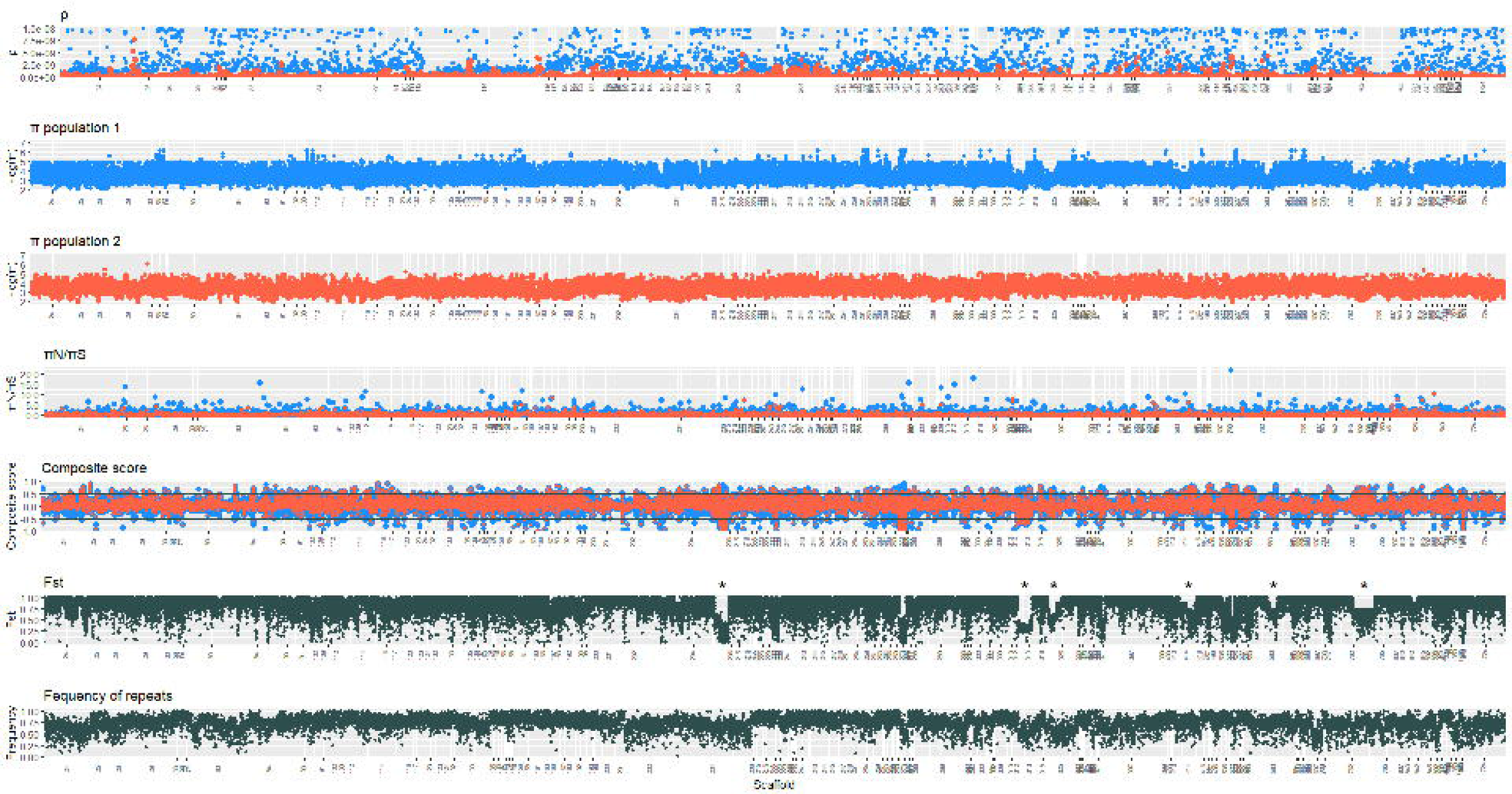

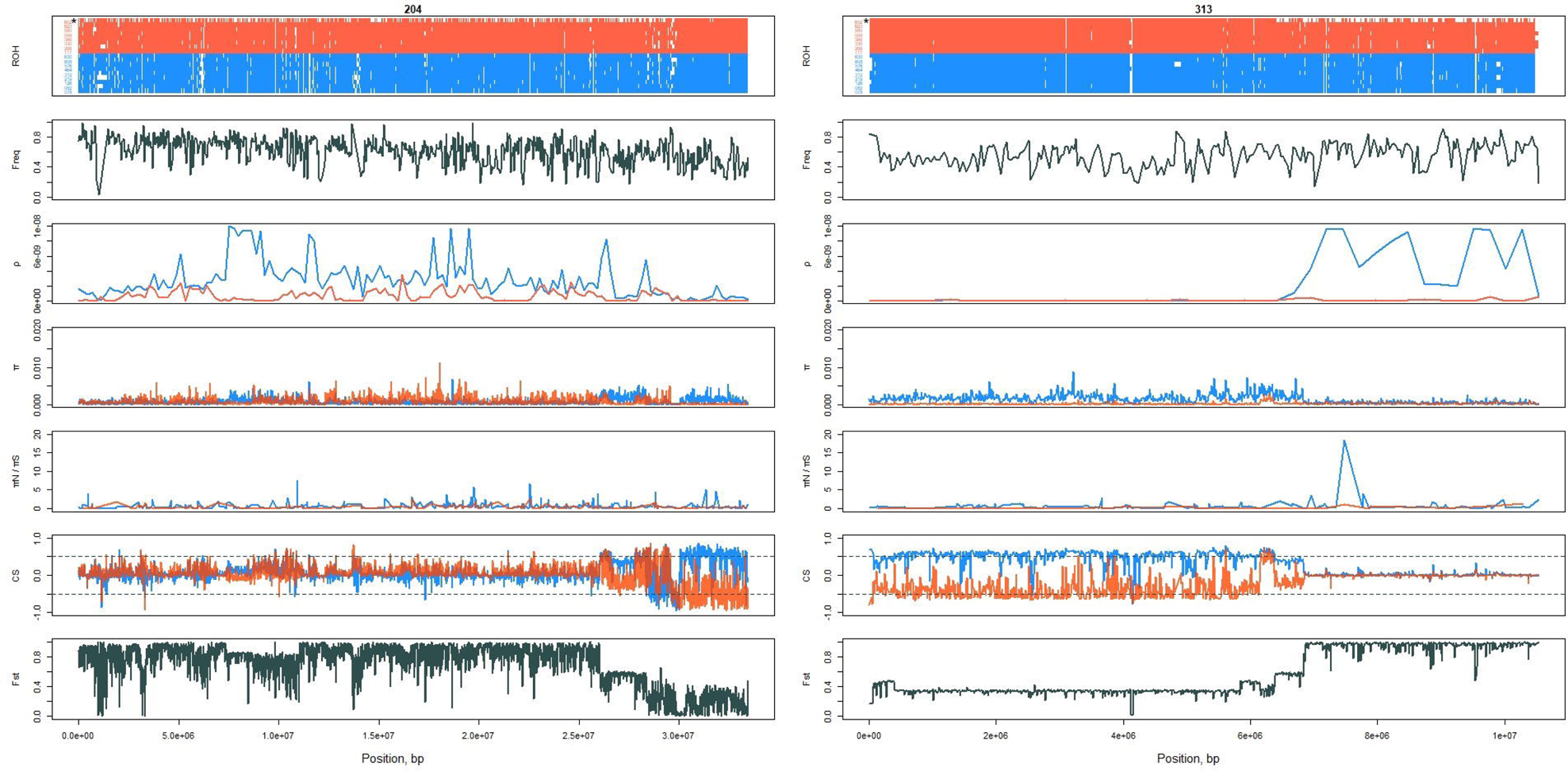

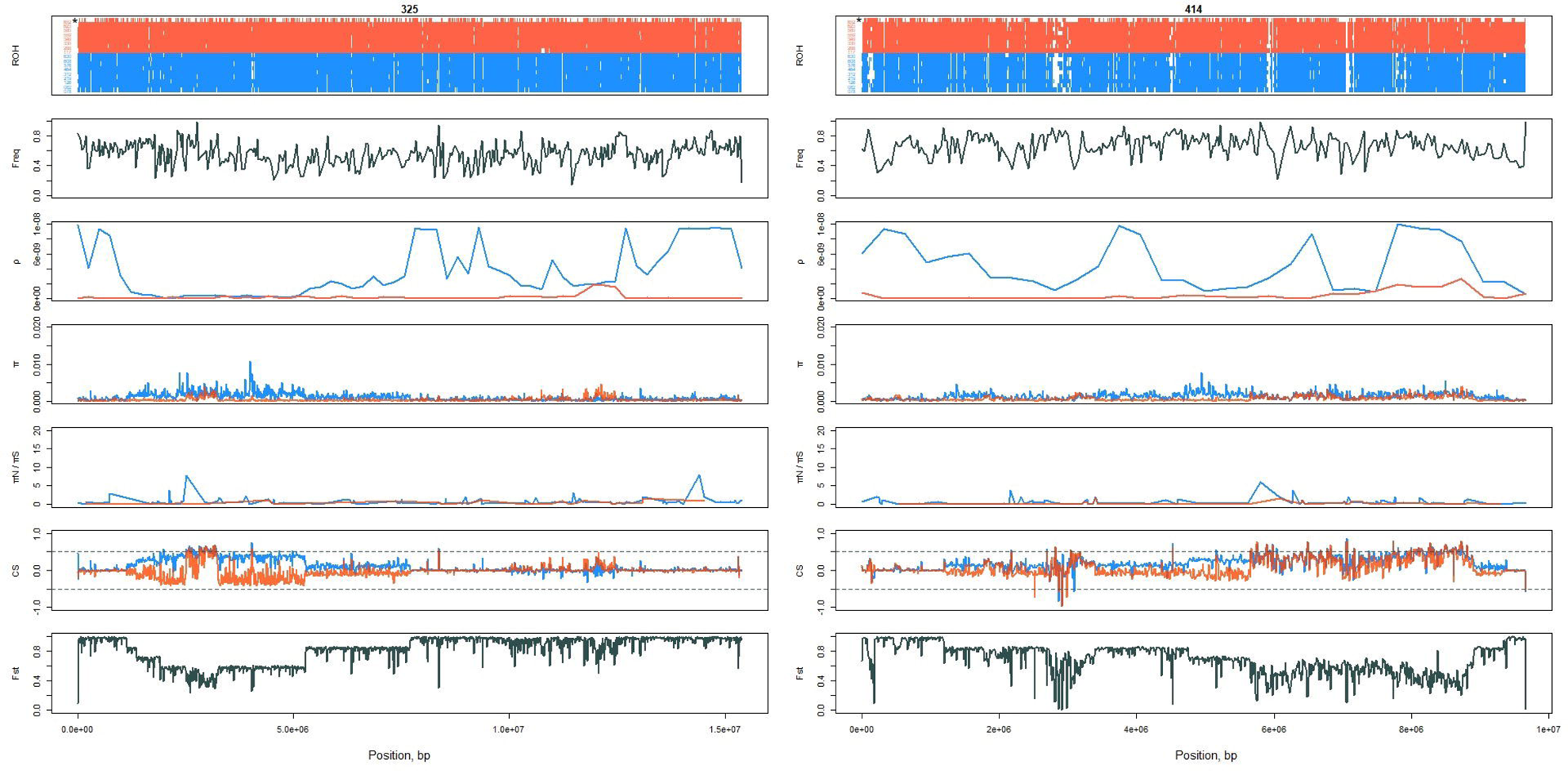

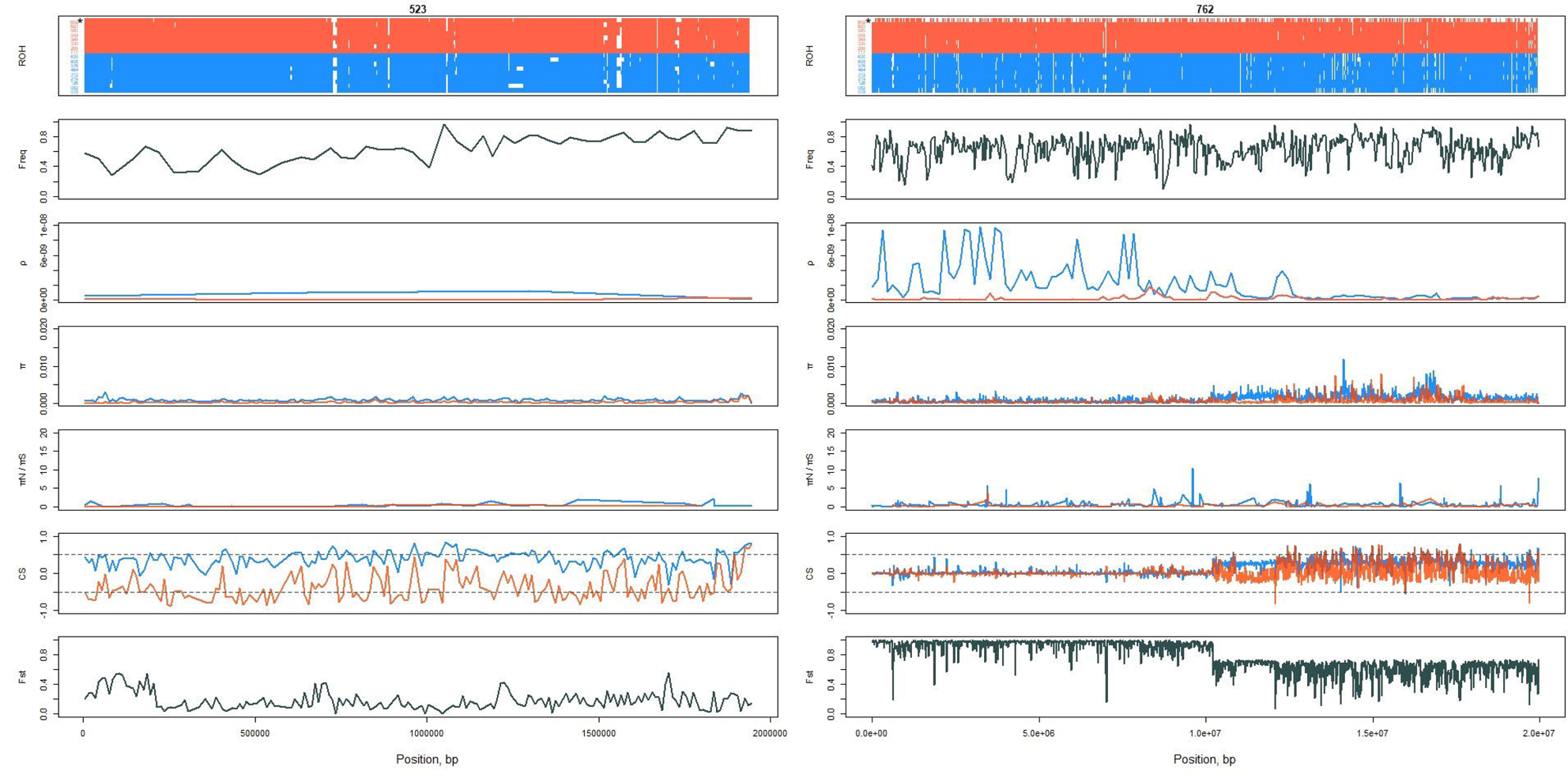

**Table 1.** Composite score outlier values detected in the *Diplolepis rosae* genome regions with low differentiation.

| Composite score (CS) outliers in *D. rosae* lineages | Scaffold number | Region, Mbp | Other signal(s) |
| --- | --- | --- | --- |
| Negative (CS < -0.5) population 1 | 204 | 30.0 | Decrease in $\rho$ |
| | 313 (**Fig. 5**) | 4.1 | Decrease in $\rho$ |
| | 414 (**Fig. 6**) | 3.0 | Decrease in $\rho$ |
| Negative (CS < -0.5) population 2 | 204 | 30.0 | Decrease in $\rho$ |
| | 204 | 30.1–33.0 | Decrease in $\rho$ |
| | 313 | 0.0–5.9 | Decrease in $\rho$ |
| | 414 | 3.0 | Decrease in $\rho$ |
|  | 523 (**Fig. 7**) | 0.0–2.0 | – |
| Positive (CS > 0.5) population 1 | 204 | 26.0 | Increase in $\rho$ |
| | 204 | 28.0 | Increase in $\rho$ |
| | 204 | 30.1–33.0 | Decrease in $\rho$ |
|  | 313 | 0.0–4.0 | – |
|  | 313 | 4.2–5.9 | – |
|  | 313 | 6.1–6.2 | – |
| | 325 (**Fig. 6**) | 2.5–3.0 | Increase in $\pi$, decrease in $\rho$, and increase in $\pi N/\pi S$ |
| | 414 | 5.5–9.0 | Increase in $\rho$ |
| | 414 | 5.8–6.0 | Increase in $\pi N/\pi S$ |
|  | 523 | 0.0–2.0 | – |
| | 762 (**Fig. 7**) | 12.5–17.5 | Increase in $\pi$ and decrease in $\rho$ |
| Positive (CS > 0.5) population 2 | 204 | 26.0 | – |
|  | 204 | 28.0 | – |
| | 313 | 6.1–6.2 | Increase in $\pi$ |
| | 325 | 2.5–3.0 | Increase in $\pi$ and decrease in $\rho$ |
|  | 414 | 5.5–9.0 | – |
| | 762 | 12.5–17.5 | Increase in $\pi$ and decrease in $\rho$ |

### *Wolbachia* identification

*Wolbachia* contigs belonging to supergroups A and B were identified in 5 and 14 *D*. *rosae* individuals, respectively (**Table S6**). In *D*. *rosae*-078 and *D*. *rosae*-117, both supergroups were identified in the same bin in the metagenome assembly. The population structure of *Wolbachia* did not follow that of *D*. *rosae* (**Fig. S12–S16**). The average number of *Wolbachia* supergroup A copies per cell (coverage) varied from 0 to 2.4. The coverage of *Wolbachia* supergroup B varied from 0 to 14.0. There was no significant difference between the *D*. *rosae* lineages in terms of *Wolbachia* coverage (supergroup A: Mann–Whitney U test, U = 25.5, z = 0.276, p = 0.782; supergroup B, U = 20, z = 0.868, p = 0.385). The ‘*Wolbachia* supergroup A/B’ ratio was the same in both *D*. *rosae* lineages (U = 23, z = 0.621, p = 0.535).

## Discussion

Our results showed that two peripatric, highly differentiated lineages of *D*. *rosae* (hereafter ‘population 1’ and ‘population 2’) occurred in France. The two lineages differed in homozygosity level and recombination rate.

Both *D*. *rosae* lineages were recovered across France, with no clear geographic distribution. Nonetheless, population 1 seemed to be more frequent in the north of France, notably in the Île-de-France region, whereas population 2 seemed to be more frequent in the southeast of France. The first structuring factor that could reflect the distribution of *D*. *rosae* is the host plant genotype, but this is unlikely to be the case. Indeed, different *D*. *rosae* genotypes have been found to parasitise the same *R*. *canina* genotype, and the same *D*. *rosae* genotype has also been observed in different *Rosa* spp. (Stille 1985; Kohnen et al. 2011). Furthermore, in our study, in three instances among 61 localities, individuals from different lineages were found on the same branch of the same host plant specimen (**Fig. 2**). Nevertheless, it is possible that each population exploits different microhabitats in *Rosa* shrubs (for example, different elevations of parasitised leaf buds or the orientation of the galls towards the sun). The second structuring factor could be an infection with different *Wolbachia* strains. However, we found that *Wolbachia* infection did not explain the population structure of *D*. *rosae*. Both lineages were infected with the same *Wolbachia* strains belonging to supergroups A and B in varying quantities. Finally, each *D*. *rosae* lineage could be under top-down control and attacked by a specific parasite community that controls population density. Local variation in the presence of parasite species would then determine the prevalence of each population. Indeed, gall wasps are exposed to intense attacks by natural enemies, notably hymenopteran parasitoids and cynipid inquilines (Stille 1984; Rizzo and Massa 2006; Todorov et al. 2012; Laszlo et al. 2014). For instance, Rizzo and Massa (2006) showed that, on average, only 5% of the *D*. *rosae* progeny per gall survived in each generation. Therefore, parasitism avoidance is under strong selection (Stone et al. 2003) and might act as a top-down structuring factor.

According to the best supported demographic scenario, the *D*. *rosae* lineages originated from a bottleneck of an ancestral population that split into two populations with no migration since the split. However, we found the same demographic scenario, with the migration parameter being equally probable as the simpler model. Indeed, we observed one admixed individual, *D*. *rosae*-652 (**Fig. 1**), belonging to population 2 but showing 13.8% of the genome assigned to population 1. This could be due to a shared ancestral polymorphism or recent gene flow. Gene flow between the two lineages is more likely because this individual showed the highest heterozygosity (**Fig. 3**) and the lowest number of widespread ROHs (**Fig. S5**). Therefore, we concluded that the two *D*. *rosae* lineages were well-differentiated populations with rare gene flow.

Background selection is expected to be a major force acting on polymorphism in thelytokous organism, such as *D*. *rosae*, because populations consist of almost only females (Nordlander 1973; Plantard et al. 1999; Stille 1985; laboratory observations) and show low levels of heterozygosity (an average of 3 and 2 heterozygous sites per 10 kbp in population 1 and population 2, respectively). A negative correlation between the density of protein-coding sequences and nucleotide diversity π usually indicates selection at linked sites, such as background selection, because deleterious alleles are more likely to occur in genes (Payseur and Nachman 2002). However, we did not observe such a pattern, which is probably the result of an intense purge of deleterious alleles (**Fig. S4**). Indeed, there are two reasons to expect the genetic load to be low in *D*. *rosae*. First, in Hymenoptera, deleterious mutations are usually purged in haploid males. Second, in thelytokous organisms, deleterious mutations are also purged in highly homozygous females (Pearcy et al. 2006).

Thelytoky also led to a reduced recombination rate and widespread ROHs, which we observed in both lineages (**Fig. 4–7**). This is concordant with previous studies suggesting that *D*. *rosae* reproduces mostly by thelytoky (Nordlander 1973; Plantard et al. 1999; Stille 1985; Stille and Dävring 1980). As previously stated by Stille and Dävring (1980) and discussed in other works (Plantard et al. 1999; Schilthuizen and Stouthamer 1998; Van Meer et al. 1995), we suggest that gamete duplication is the main mechanism of thelytoky in *D*. *rosae*. Indeed, gamete duplication leads to complete homozygosity across the genome. Although we did not observe complete homozygosity, we detected ROHs reaching several Mbp and making up to 90% of the genome, which is concordant with frequent gamete duplication thelytoky. Thus, other mechanisms of thelytoky retaining local heterozygosity (Rabeling and Kronauer 2013), e.g. via central or terminal fusion or apomixis, are less likely to occur or less frequent. Another clue that *D*. *rosae* reproduces by gamete duplication thelytoky is the presence of *Wolbachia*. *Wolbachia* is known to induce thelytoky via gamete duplication in Hymenoptera (Stouthamer et al. 1990; Stouthamer and Kazmer 1994; Gottlieb et al. 2002). It has also been suggested to be the case in other *Diplolepis* sp. (Plantard et al. 1998). However, we acknowledge that we have only indirect proof of gamete duplication thelytoky. Only laboratory experiments should be able to provide clear-cut evidence (for example, Stouthamer et al. 1990).

Both thelytokous *D*. *rosae* lineages differed in the intensity of recombination and heterozygosity. Population 1 showed higher heterozygosity and higher recombination rates than population 2. This could reflect sexual reproduction or the use of an alternative mechanism of thelytoky. To reveal the factors that explain the difference between the two lineages, we scanned the *D*. *rosae* genome for regions deviating from the background. Using the composite score (CS) summarising Fst and π values, we unravelled regions with low differentiation and low or high nucleotide diversity. Indeed, the decrease in Fst indicates low differentiation between lineages, which is maintained by local gene flow or by selection. If it is associated with low diversity, it could indicate negative selection on linked sites that have maintained low differentiation between lineages since their split. If it is associated with high diversity, it could reflect balancing selection when the same diversity is maintained since the population split. It could also be a relaxed negative selection that allows a new allele to emerge that would have been previously deleterious. We acknowledge that this approach has some limitations in detecting selection. First, it is challenging to distinguish between balancing selection/relaxed constraint and the effect of recombination in the regions showing a local increase in nucleotide diversity. Therefore, we estimated the recombination rate across the genome to contrast the diversity level and recombination. Therefore, when we observed a high composite score in the non-recombining regions, putative balancing selection or relaxed constraint was likely at play. Second, the method is limited in terms of negative and balancing selection. In the case of positive selection or negative selection acting on different haplotypes in the different lineages, we observed high Fst, low π, and CS close to 0. However, the Fst between the two lineages was already close to 1 in most of the genome (**Fig. 4**). Therefore, to properly detect positive selection, one should perform other analyses (for example, the McDonald–Kreitman test that needs an outgroup) that are not affected by the mode of reproduction or demography (McDonald and Kreitman 1991; Eyre-Walker 2002; Parsch et al. 2009).

Despite these limitations, we revealed that the two lineages demonstrated opposite composite score outliers in several ROHs, which could be due to contrasted selective regimes. In the more recombining population 1, the genes associated with ‘sperm competition’, ‘insemination’, ‘copulation’, ‘metallopeptidase activity’, and ‘metalloendopeptidase activity’ terms showed patterns typical of negative selection, while in the less recombining population 2, they would be putatively under balancing or relaxed selection. Genes associated with the term ‘commissural neuron axon guidance’ showed the opposite trend with genomic signatures of balancing/relaxed selection in population 1 and negative selection in population 2. The changes in the selective regime of those specific genes could be related to different selective processes acting in the *D*. *rosae* lineages and could explain the maintenance of the two populations despite a shared habitat. In population 1, genes involved in sexual reproduction are important for efficient recombination through the production of males, thereby generating genetic diversity in terms of allele combination. Higher genetic diversity would then give an advantage in survival in the face of a highly prevalent and diverse community of parasites (Stille 1984; Rizzo and Massa 2006; Todorov et al. 2012; Laszlo et al. 2014). Our suggestion could be supported by the study of Rizzo and Massa (2006), which showed a possible association between average parasitism and the percentage of *D*. *rosae* males that emerged. Parasitism rates were 30.5% and 34.3% in two *D*. *rosae* populations consisting of 21% and 15.6% males, respectively. In other populations, there were no males, and the average parasitism rate reached 57.6% (varied from 12.5% to 100%) (Rizzo and Massa 2006). In population 2, showing more widespread ROHs and lower recombination, the candidate genes from the terms ‘sperm competition’, ‘insemination’, and ‘copulation’ showed signatures of either balancing selection or relaxed selection. Regarding the extreme level of homozygosity in population 2, male-related genes could become unnecessary and be under relaxed selection. However, balancing selection could also take place to maintain male alleles through frequency-dependent selection, as alleles important for male function would increase in frequency when selection for recombination occurs. In contrast to male-related function, the genes involved in ‘commissural neuron axon guidance’ showed a signature typical of purifying selection in population 2 but balancing selection or relaxed constraint in population 1. We hypothesise that a less recombining lineage shows a conservative host-searching behavioural pattern. Indeed, Ramirez-Romero et al. (2012) demonstrated differences in host-searching behaviour between thelytokous and arrhenotokous populations of *Odontosema anastrephae* (Hymenoptera: Cynipidae s. lat.), the figitid parasitoid of the fruit fly. In this study, thelytokous females showed a basic behavioural sequence when exploring the odour source, whereas arrhenotokous females demonstrated more complex behaviour (Ramirez-Romero et al. 2012).

## Conclusion

We demonstrated the existence of two highly differentiated peripatric lineages of *D*. *rosae* that differ in the level of recombination and homozygosity across the genome. The maintenance of these populations might be due to selection acting upon different traits. Further research could explore the natural history of the two populations by examining the following questions. Does each lineage really differ in the frequency of male production? Are there differences in the parasite communities attacking them that would explain top-down control? Do they show the same phenology? These are key questions to answer to determine which selective pressure leads to the maintenance of recombination in *D*. *rosae*, despite mostly thelytokous reproduction.

## Materials and Methods

### Sampling

Eighteen dog rose (*Rosa canina*) bud galls with *D*. *rosae* were collected from April 2020 to December 2020 in France for sequencing (**Table S7**). After sampling, the galls were kept in plastic bags at room temperature. The insect material was removed by dissection of the gall tissue for further DNA extraction. A one-tube sample contained emerged adults or larvae from the same gall. Further DNA extractions were performed from larvae or adult females (**Table S7**).

### DNA extraction for Illumina sequencing

Prior to DNA extraction, the insects were frozen at −80°C overnight. The initial mass of the material provided for DNA extraction varied from 13.7 to 56.0 mg. The insect material was homogenised using a TissueLyser (Qiagen, Haan, Germany) with a metallic bead. Insect DNA was extracted using a DNeasy Blood and Tissue Kit (Qiagen, Germany). After extraction, the purity of the samples was evaluated by estimating the 260/280 and 260/230 ratios measured using a Thermo Scientific NanoDrop 2000 Spectrophotometer. The 260/280 ratio varied from 1.69 to 2.08; the 260/230 ratio varied from 0.79 to 2.08. The DNA quantity was estimated using a Qubit dsDNA BR Assay Kit (Invitrogen) and the Qubit Fluorometer and varied from 0.98 to 6.9 µg. Illumina sequencing was performed by Genotul, Toulouse, France (https://get.genotoul.fr). The Illumina samples are available in the NCBI database under the BioSample accessions from SAMN32903234 to SAMN32903250.

### DNA extraction for nanopore sequencing

Prior to DNA extraction, one *D*. *rosae* larva (**Table S6**) was ground by hand with a sterile pestle. The DNA was extracted using the NucleoBond Buffer Set IV kit (Macherey-Nagel, Germany) and NucleoBond AXG 20 columns (Macherey-Nagel, Germany). Long-read sequencing was performed using a Flow Cell Wall Wash Kit (EXP-WSH004) (Oxford Nanopore Technologies, UK) and an Oxford Nanopore MinION Flow Cell R10 (Oxford Nanopore Technologies, UK).

### Genome assembly

Oxford Nanopore Technology (ONT) raw reads were base called using Guppy basecaller v. 6.1.2+e0556ff (Oxford Nanopore Technologies, UK) with accurate mode and dna_r9.4.1_450bps_sup.cfg config. The resulting .*fastq* reads were assembled with a Flye assembler (Kolmogorov et al. 2019) using a --nano-hq flag, an estimated coverage of 18x, and an error rate of 0.1. The input genome size was 580 Mbp, based on a k-mer estimate using GenomeScope (Vurture et al. 2017). Because of the high error rate of ONT reads, the resulting assembly was polished using NextPolish (Hu et al. 2020) with high-coverage Illumina reads (119x) using a sample from the same population (sample ESE-219, **Table S6**). Repeats were discovered in the de novo assembly using RepeatModeler v. 2.0.3 (Flynn et al. 2020) using RMBlast v. 2.11.0. *De novo*-detected repeats were assessed for potential wrong assignment to repeats of highly duplicated genes by blasting each consensus repeat sequence onto the NCBI nr database. Repeats matching known protein-coding genes not related to known repeats were removed, and filtered repeats were then used to build the repeat database. *De novo* assembly was then masked using the newly built repeat database. The frequency of repeats was calculated in sliding windows and presented as the ratio between the repeat length and the length of the given genome region. The assembly quality was evaluated by computing different metrics using QUAST v. 5.1.0rc1 (Mikheenko et al. 2018). Protein-coding genes were predicted using BRAKER2 with ab-initio mode (Hoff et al. 2019) using protein homology from OrthoDB Metazoa, Fungi, Plants, and Bacteria for the ProtHint (Bruna et al. 2020) step. The completeness of annotated genes in terms of their expected gene content was evaluated using BUSCO v. 5.3.2 (Manni et al. 2021). Predicted genes were functionally annotated using the eggNOG-mapper v. 2.1.7 web server (Huerta-Cepas et al. 2019; Cantalapiedra et al. 2021).

### SNP calling

All reads of *D*. *rosae* were first aligned to the reference using Bowtie 2 v. 2.3.5.1 (Langmead and Salzberg 2012). The alignments were processed for further manipulation using samtools v. 1.7 (view, sort, index, depth, and stat options) (Danecek et al. 2021). The quality of each alignment was estimated by calculating the average percentage of the mapped reads (57.3–98.1%), the average quality (35.4–35.7), the average coverage (25–119×), and the error rate (0.0078–0.015). Subsequently, the identification of polymorphisms across the genome was performed using a pipeline described in GenomeAnalysisToolkit (GATK) v. 4.0 (AddOrReplaceReadGroups, HaplotypeCaller, CombineGVCFs, and GenotypeGVCFs tools) (https://gatk.broadinstitute.org/hc/en-us/articles/360035890411-Calling-variants-on-cohorts-of-samples-using-the-HaplotypeCaller-in-GVCF-mode). Indels and non-biallelic sites were removed from the .*vcf* file using bcftools v. 1.7 (filter command) (Danecek et al. 2021). The rare (--maf 0.05), low-quality (--minQ 30), low-depth (cutoff below 2.5th percentile), and high-depth (cutoff above 97.5th percentile) sites were removed using vcftools 0.1.17 (Danecek et al. 2021). The .*vcf* file was phased by beagle v. 5.4 (Browning et al. 2021). The total number of examined polymorphisms was 2,903,839.

### Population structure

The population structure of *D*. *rosae* was inferred according to the fastStructure algorithm (Raj et al. 2014) based on the calculation of the allele frequency spectra from SNP data. Prior to fastStructure, indels and site linkage disequilibrium (r^2^>0.2) were filtered out using bcftools v. 1.7 (filer and prune commands) (Danecek et al. 2021) from the initial .*vcf* file that was then converted to a .*bed* file by plink v. 1.9 (Purcell et al. 2007). After filtering, the total number of examined SNPs was 12,309. The population structure was assessed with the number of populations, K, varying from 1 to 4. To show individuals admixed between the lineages, F3 statistics representing the covariance of allele frequency differences between populations were calculated using ADMIXTOOLS 2.0.0 (Patterson et al. 2012; Peter 2016).

### DNA extraction for genotyping

Prior to DNA extraction, the insects were frozen at −20°C overnight. The DNA was extracted using either a DNeasy Blood and Tissue Kit (Qiagen, Germany) or Chelex 100 Resin (Bio-Rad).

### Genotyping

A genetic marker helping to distinguish the lineages of *D*. *rosae* was searched along the genome by choosing a 15 kbp window showing the Fst value closest to 1 and substantial polymorphism. A 706-bp sequence containing 9 SNPs was selected, and the primers were designed using Primer3web 4.1.0 (Untergasser et al. 2012). The marker was then amplified using polymerase chain reaction (PCR) (**Protocol S1**). The presence of the PCR product was verified by performing electrophoresis in 3.5% agarose gel and sequencing using Eurofins Genomics. The sequences were assessed visually on the trace file using SnapGene Viewer 6.0.2 (“SnapGene software” www.snapgene.com) and quality trimmed. Subsequently, the sequences were used for cladogram construction (maximum likelihood statistical method, Tamura-Nei substitution model (default)) by MEGA 11 (Tamura et al. 2021). A total of 123 *D*. *rosae* individuals from 61 locations were genotyped for the marker. Samples were collected from different habitats in France, from different host plant individuals from the same habitat, and from different galls from the same host plant.

### Demographic scenarios

Scenarios describing the demographic history of *D*. *rosae* were examined using *dadi* software based on the diffusion (continuous approximation) approach (Gutenkunst et al. 2009). The following standard two-population models were examined: bottleneck followed by exponential growth, then split without (i) and with migration (ii), split into two populations of specified size (iii), isolation with exponential population growth (iv), isolation with exponential population growth, and a size change prior to splitting (v) (**Code S1**). A set of joint allele frequency spectra (AFS) was generated to compare the model with the data. Each model was examined by varying population sizes, time of split/isolation, and migration parameter (0, symmetric, or asymmetric). The models were ranked by calculating the Akaike Information Criterion (AIC). A likelihood ratio test was performed to compare the best models according to AIC.

### Estimation of model parameters

To confirm the demographic scenario found by *dadi* and estimate parameters describing the demography of *D*. *rosae* (**Code S2**), the approximate Bayesian computation (ABC) was applied. One million simulations of the population parameters (effective population size Ne, mutation rate μ, bottleneck time, split time) were performed, and the resulting summary statistics were calculated by the msprime simulator v. 1.1.1 (Baumdicker et al. 2022). Subsequently, the goodness-of-fit and validation of the model were performed using the *abc* R package v. 2.2.1 (Csilléry et al. 2012).

### Population genomic statistics

Absolute divergence D_xy_ (Nei 1987) and the fixation index Fst (Weir and Cockerham 1984) were calculated in the genome 10-kbp windows using pixy v. 1.2.6.beta1 (Korunes and Samuk 2021). Tajima’s D and nucleotide diversity π (in 10-kbp windows) and per-individual heterozygosity H were calculated using vcftools 0.1.17 (Danecek et al. 2021). The Watterson estimator θ*w* was calculated as the number of segregating sites (provided in the output table using vcftools 0.1.17 when calculating π) in the genome 10-kbp windows divided by the sum of the (n−1) first harmonic means, where n is the number of haplotypes (Watterson 1975). Nucleotide diversity at nonsynonymous π*_N_* and synonymous π_S_ sites was calculated in protein coding sequences (CDS) using the PolydNdS programme (Thornton 2003). Each CDS was extracted from the genome using the vcf2fasta.py script (https://github.com/santiagosnchez/vcf2fasta). The effect of background selection on the genome was assessed by measuring the correlation between gene density (the proportion of nucleotides assigned to a protein-coding sequence) and π within each scaffold. The population-scaled recombination rate ρ was estimated by ReLERNN (Adrion et al. 2020) using the ReLERNN_SIMULATE - ReLERNN_TRAIN - ReLERNN_PREDICT - ReLERNN_BSCORRECT pipeline. In ReLERNN_TRAIN, -- nEpochs (time) was estimated at 500, corresponding to a minimal convergence of loss (mean squared error) between the training set and the validation set.

### Runs of homozygosity

ROHs were detected across the *D*. *rosae* genome using bcftools v. 1.7 (roh command) with the option -G, the phred-scaled genotype likelihoods, set to 30 (Danecek et al. 2021). The frequency of 0.01–0.1, 0.1–0.5, 0.5–1.0, and >1 Mbp runs was estimated as the total number of ROHs from each category divided by the total number of ROHs.

### Detection of genome regions under selection

To detect regions under putative selection in the *D*. *rosae* genome, we calculated Fst and π in 10-kbp windows using pixy v. 1.2.6.beta1 (Korunes and Samuk 2021). To distinguish between genomic regions with low differentiation/high nucleotide diversity and low differentiation/low diversity, we created a composite score that summarised Fst and π. The score was equal to (1 − Fst)*2(F(π) − 0.5), where F(π) is the cumulative distribution function in each lineage. Composite score outliers below −0.5 reflected genomic regions with low differentiation and low genetic diversity, while composite score outliers above 0.5 indicated regions with low differentiation and high diversity (**Fig. S14, Fig. S15**). Subsequently, annotated gene sets (eggNOG-mapper) found in these regions were used in the Gene set enrichment analysis (GSEA). GSEA was performed using the *topGO* R package v. 2.48.0 by applying the Fisher exact test (Alexa and Rahnenfuhrer 2022). The obtained raw p-values were adjusted using the *p*.*adjust* function (R Core Team 2022; https://www.R-project.org/). In total, 8721 genes were used. The examined number of genes showing a composite score below −0.5 was 75 for population 1 and 333 for population 2, respectively. The examined number of genes showing a composite score above 0.5 was 177 and 80 for population 1 and population 2, respectively.

### Identification of *Wolbachia*

The assembly of the *D*. *rosae* genome using Nanopore data shows only one completely assembled genome of *Wolbachia* belonging to supergroup B (Wang et al. 2020). To distinguish between *Wolbachia* genomes from supergroup B and supergroup A, contaminant reads were removed from the *D*. *rosae* reference (Bowtie 2 --un-conc-gz option) and assembled using the MEGAHIT metagenome assembler v. 1.2.9 (Li et al. 2015). The obtained contigs were used to reconstruct genomes with MetaBAT v. 2.12.1 (Kang et al. 2015). After binning, the *Wolbachia* supergroups were identified according to the multilocus sequence typing (MLST) system based on five genes (*coxA*, *gatB*, *hcpA*, *ftsZ*, and *fbpA*) (Wang et al. 2020). The coverage of *Wolbachia* contigs was given by MetaBAT and normalised by the coverage of the corresponding *D*. *rosae* individual.

### Statistics

All statistical analyses were performed using R v 4.2.2 (R Core Team 2022; https://www.R-project.org/). The significance level was set to 0.05. The figures were produced using R v. 4.2.2 and Microsoft Excel 2010.

## Acknowledgements

We would like to thank David Ogereau (CNRS, EGCE) for providing DNA for Nanopore sequencing. We also thank Jacqui Shykoff (Paris-Saclay University, ESE), Olivier Plantard (INRAE), Amir Yassin (CNRS, EGCE), Florence Mougel (CNRS, EGCE), and Thierry Robert (Paris-Saclay University, ESE). The study received a grant from the National Agency for Research (France) (Project ANR-19-CE02-0008 “Tracing back the history of an adaptive trait: genetic basis of plant host manipulation by gall wasps - BETAGALL”).

## Author Contributions

KM performed DNA extraction (genotyping), simulations, and data analysis, and wrote the manuscript. AB conceived and designed the study, wrote the manuscript, performed genome assembly, described the *D*. *rosae* genome, and performed GSEA. AB, KM, and ZT collected gall samples. ZT kept the gall samples and performed DNA extraction (Illumina). All authors discussed the results and contributed to the final manuscript.

## Data Availability

The assembled genome of *D*. *rosae* is available at DDBJ/ENA/GenBank under accession JAPYXD000000000. The Illumina *D*. *rosae* reads are available in the NCBI database under BioSample accessions from SAMN32903234 to SAMN32903250.

## Literature cited

Adachi-Hagimori T, Miura K, Abe Y. 2011. Gene flow between sexual and asexual strains of parasitic wasps: a possible case of sympatric speciation caused by a parthenogenesisLinducing bacterium. J Evol Biol. 24(6):1254–1262.

Adrion JR, Galloway JG, Kern AD. 2020. Predicting the landscape of recombination using deep learning. Mol Biol Evol. 37(6):1790–1808.

Alexa A, Rahnenfuhrer J. 2022. topGO: Enrichment Analysis for Gene Ontology. R package version 2.50.0. doi: 10.18129/B9.bioc.topGO

Baumdicker F, et al. 2022. Efficient ancestry and mutation simulation with msprime 1.0. Genetics. 220(3):iyab229.

Belshaw R, Quicke DL. 2003. The cytogenetics of thelytoky in a predominantly asexual parasitoid wasp with covert sex. Genome. 46(1):170–173.

Bordenstein SR. 2003. 17 symbiosis and the origin of species. Insect Symbiosis:283 –304.

Browning BL, Tian X, Zhou Y, Browning SR. 2021. Fast two-stage phasing of large-scale sequence data. Am J Hum Genet. 108(10):1880–1890.

Bruna T, Lomsadze A, Borodovsky M. 2020. GeneMark-EP and-EP+: automatic eukaryotic gene prediction supported by spliced aligned proteins. bioRxiv. 17. doi: 10.1101/2019.12.31.891218.

Cantalapiedra CP, Hernández-Plaza A, Letunic I, Bork P, Huerta-Cepas J. 2021. eggNOG-mapper v2: functional annotation, orthology assignments, and domain prediction at the metagenomic scale. Mol Biol Evol. 38(12):5825–5829.

Capdevielle-Dulac C, et al. 2022. Spontaneous parthenogenesis in the parasitoid wasp *Cotesia typhae*: low-frequency anomaly or evolving process? Peer Commun J. 2: e37.

Csilléry, K., François, O., & Blum, M. G. 2012. abc: an R package for approximate Bayesian computation (ABC). Methods Ecol Evol. 3(3):475–479.

Crow JF, Kimura M. 1965. Evolution in sexual and asexual populations. Am Nat. 99(909):439–450.

Danecek P, et al. 2021. Twelve years of SAMtools and BCFtools. GigaScience. 10(2):giab008.

Engelstädter J, Sandrock C, Vorburger C. 2011. Contagious parthenogenesis, automixis, and a sex determination meltdown. Evol: Int J Org Evol. 65(2):501–511.

Eyre-Walker A. 2002. Changing effective population size and the McDonald-Kreitman test. Genetics. 162(4):2017–2024.

Flynn JM, et al. 2020. RepeatModeler2 for automated genomic discovery of transposable element families. Proc Natl Acad Sci USA. 117(17):9451–9457.

Foray V, Henri H, Martinez S, Gibert P, Desouhant E. 2013. Occurrence of arrhenotoky and thelytoky in a parasitic wasp *Venturia canescens* (Hymenoptera: Ichneumonidae): effect of endosymbionts or existence of two distinct reproductive modes?. Eur J Entomol. 110(1):103–107.

Giorgini M, Monti MM, Caprio E, Stouthamer R, Hunter MS. 2009. Feminization and the collapse of haplodiploidy in an asexual parasitoid wasp harboring the bacterial symbiont *Cardinium*. Heredity. 102(4):365–371.

Giron D, Huguet E, Stone GN, Body M. 2016. Insect-induced effects on plants and possible effectors used by galling and leaf-mining insects to manipulate their host-plant. J Insect Physiol. 84:70–89.

Gottlieb Y, Zchori-Fein E, Werren JH, Karr TL. 2002. Diploidy restoration in *Wolbachia*-infected *Muscidifurax uniraptor* (Hymenoptera: Pteromalidae). J Invertebr Pathol. 81(3):166–174.

Gutenkunst RN, Hernandez RD, Williamson SH, Bustamante CD. 2009. Inferring the joint demographic history of multiple populations from multidimensional SNP frequency data. PLoS Genetics. 5(10):e1000695.

Hamilton WD. 1980. Sex versus non-sex versus parasite. Oikos. 35:282–290.

Heimpel GE, De Boer JG. 2008. Sex determination in the Hymenoptera. Annu Rev Entomol. 53:209–230.

Hoff KJ, Lomsadze A, Borodovsky M, Stanke M. 2019. Whole-genome annotation with BRAKER. Methods Mol Biol. 1962:65–95. 10.1007/978-1-4939-9173-0_5.

Hu J, Fan J, Sun Z, Liu S. 2020. NextPolish: a fast and efficient genome polishing tool for long-read assembly. Bioinformatics. 36(7):2253–2255.

Huerta-Cepas J, et al. 2019. eggNOG 5.0: a hierarchical, functionally and phylogenetically annotated orthology resource based on 5090 organisms and 2502 viruses. Nucleic Acids Res. 47(D1):D309–D314.

Kang DD, Froula J, Egan R, Wang Z. 2015. MetaBAT, an efficient tool for accurately reconstructing single genomes from complex microbial communities. PeerJ. 3:e1165.

Kohnen A, Wissemann V, Brandl R. 2011. No host-associated differentiation in the gall wasp *Diplolepis rosae* (Hymenoptera: Cynipidae) on three dog rose species. Biol J Linn Soc. 102(2):369–377.

Kolmogorov M, Yuan J, Lin Y, Pevzner PA. 2019. Assembly of long, error-prone reads using repeat graphs. Nat Biotechnol. 37(5):540–546.

Korunes KL, Samuk K. 2021. pixy: unbiased estimation of nucleotide diversity and divergence in the presence of missing data. Mol Ecol Resour. 21(4):1359–1368.

Kozek WJ & Rao RU. 2007. The discovery of Wolbachia in arthropods and nematodes – a Historical perspective. In: Wolbachia: A bug’s life in another bug. Issues in Infectious Diseases. Vol. 5. pp. 1–14.

Langmead B, Salzberg SL. 2012. Fast gapped-read alignment with Bowtie 2. Nat Methods. 9(4):357–359.

Laszlo Z, Solyom K, Prazsmari H, Barta Z, Tothmeresz B. 2014. Predation on rose galls parasitoids and predators determine gall size through directional selection. PLoS One 9(6):e99806.

Li D, Liu CM, Luo R, Sadakane K, Lam TW. 2015. MEGAHIT: an ultra-fast single-node solution for large and complex metagenomics assembly via succinct de Bruijn graph. Bioinformatics. 31(10):1674–1676.

Manni M, Berkeley MR, Seppey M, Zdobnov EM. 2021. BUSCO: assessing genomic data quality and beyond. Curr Protocol. 1:e323.

Maynard-Smith J. 1978. The evolution of sex. Cambridge: Cambridge University Press.

McDonald JH, Kreitman M. 1991. Adaptive protein evolution at the *Adh* locus in *Drosophila*. Nature. 351(6328):652–654.

Mikheenko A, Prjibelski A, Saveliev V, Antipov D, Gurevich A. 2018. Versatile genome assembly evaluation with QUAST-LG. Bioinformatics. 34:i142–i150.

Nei M. 1987. Molecular evolutionary genetics. New York Chichester, West Sussex, Columbia University Press.

Nordlander G. 1973. Parasitsteklar i galler av *Diplolepis rosae* (L.) och *D. mayri* Schlechtd. (Hym. Cynipidae) (Hym. Ichneumonoidea, Chalcidoidea, Cynipoidea). Entomologisk Tidskrift. 94(3–4):148–176.

Parsch J, Zhang Z, Baines JF. 2009. The influence of demography and weak selection on the McDonald–Kreitman test: an empirical study in *Drosophila*. Mol Biol Evol. 26(3):691–698.

Patterson N, et al. 2012. Ancient admixture in human history. Genetics. 192(3):1065–1093.

Payseur BA, Nachman MW. 2002. Gene density and human nucleotide polymorphism. Mol Biol Evol. 19(3):336–340.

Pearcy M, Hardy O, Aron S. 2006. Thelytokous parthenogenesis and its consequences on inbreeding in an ant. Heredity. 96(5):377–382.

Peter BM. (2016). Admixture, population structure, and F-statistics. Genetics. 202(4):1485–1501.

Plantard O, Rasplus JY, Mondor G, Clainche IL, Solignac M. 1998. *Wolbachia*–induced thelytoky in the rose gall wasp *Diplolepis spinosissimae* (Giraud) (Hymenoptera: Cynipidae), and its consequences on the genetic structure of its host. Proc R Soc Ser B Biol Sci. 265(1401):1075–1080.

Plantard O, Rasplus JY, Mondor G, Le Clainche I, Solignac M. 1999. Distribution and phylogeny of *Wolbachia* inducing thelytoky in Rhoditini and ‘Aylacini’ (Hymenoptera: Cynipidae). Insect Mol Biol. 8(2):185–191.

Purcell S, et al. 2007. PLINK: a tool set for whole-genome association and population-based linkage analyses. Am J Hum Genet. 81(3):559–575.

Queffelec J, Allison JD, Greeff JM, Slippers B. 2021. Influence of reproductive biology on establishment capacity in introduced Hymenoptera species. Biol Invasions. 23(2):387–406.

R Core Team. 2022. R: A Language and Environment for Statistical Computing. Vienna, Austria: R Foundation for Statistical Computing. https://www.R-project.org/.

Rabeling C, Kronauer DJ. 2013. Thelytokous parthenogenesis in eusocial Hymenoptera. Annu Rev Entomol. 58:273–292.

Raj A, Stephens M, Pritchard JK. 2014. fastSTRUCTURE: variational inference of population structure in large SNP data sets. Genetics. 197(2):573–589.

Ramirez-Romero R, Sivinski J, Copeland CS, Aluja M. 2012. Are individuals from thelytokous and arrhenotokous populations equally adept as biocontrol agents? Orientation and host searching behavior of a fruit fly parasitoid. BioControl. 57(3):427–440.

Rizzo MC, Massa B. 2006. Parasitism and sex ratio of the Bedeguar gall wasp *Diplolepis rosae* (L.) (Hymenoptera: Cynipidae) in Sicily (Italy). J Hymen Res. 15(2):2779285.

Schilder K, Heinze J, Gross R, Hölldobler B. 1999. Microsatellites reveal clonal structure of populations of the thelytokous ant *Platythyrea punctata* (F. Smith) (Hymenoptera; Formicidae). Mol Ecol. 8(9):1497–1507.

Schilthuizen M, Stouthamer R. 1998. Distribution of *Wolbachia* among the guild associated with the parthenogenetic gall wasp *Diplolepis rosae*. Heredity. 81(3):270–274.

Schön I, Martens K, van Dijk P. 2009. Lost Sex. The Evolutionary Biology of Parthenogenesis. Springer.

Shorthouse JD, Floate KD. 2010. Galls induced by cynipid wasps of the genus *Diplolepi* (Hymenoptera: Cynipidae) on the roses of Canada’s grasslands. In: Arthropods of Canadian Grasslands: Ecology and Interactions in Grassland Habitats. Biological Survey of Canada, Ottawa, Ontario, Canada. pp. 251–279.

Stenberg P, Saura A. 2009. Cytology of asexual animals. In: Lost Sex. Dordrecht: Springer. p. 63–74.

Stille B. 1984. The effect of hostplant and parasitoids on the reproductive success of the parthenogenetic gall wasp *Diplolepis rosae* (Hymenoptera, Cynipidae). Oecologia. 63(3):364–369.

Stille B. 1985. Population genetics of the parthenogenetic gall wasp *Diplolepis rosae* (Hymenoptera, Cynipidae). Genetica. 67(2):145–151.

Stille BO, Dävring L. 1980. Meiosis and reproductive strategy in the parthenogenetic gall wasp *Diplolepis rosae* (L.) (Hymenoptera, Cynipidae). Hereditas. 92(2):353–362.

Stone GN, Graham N, Schönrogge K. 2003. The adaptive significance of insect gall morphology. Trends Ecol Evol. 18(10):512–522.

Stouthamer R, Kazmer DJ. 1994. Cytogenetics of microbe-associated parthenogenesis and its consequences for gene flow in *Trichogramma* wasps. Heredity. 73(3):317–327.

Stouthamer R, Luck RF, Hamilton WD. 1990. Antibiotics cause parthenogenetic *Trichogramma* (Hymenoptera/Trichogrammatidae) to revert to sex. Proc Natl Acad Sci USA. 87(7):2424–2427.

Tamura K, Stecher G, Kumar S. 2021. MEGA11: molecular evolutionary genetics analysis version 11. Mol Biol Evol. 38(7):3022–3027.

Thornton K. 2003. Libsequence: a C++ class library for evolutionary genetic analysis. Bioinformatics. 19(17):2325–2327.

Todorov I, Stojanova A, Parvanov D, Boyadzhiev P. 2012. Studies on the gall community of *Diplolepis rosae* (Hymenoptera: Cynipidae) in Vitosha Mountain, Bulgaria. Acta Zool Bulg. 4:27–37.

Tvedte ES, Logsdon Jr JM, Forbes AA. 2019. Sex loss in insects: causes of asexuality and consequences for genomes. Curr Opin Insect Sci. 31:77–83.

Untergasser A, et al. 2012. Primer3—new capabilities and interfaces. Nucleic Acids Res. 40(15):e115.

Van Meer M, van Kan FJPM, Breeuwer JAJ, Stouthamer R. 1995. Identification of symbionts associated with parthenogenesis in *Encarsia formosa* (Hymenoptera: Aphelinidae) and *Diplolepis rosae* (Hymenoptera. Cynipidae). Proc Exp Appl Entomol. 6:1–86.

Vurture GW, et al. 2017. GenomeScope: fast reference-free genome profiling from short reads. Bioinformatics. 33(14):2202–2204.

Wang X, et al. 2020. Phylogenomic analysis of *Wolbachia* strains reveals patterns of genome evolution and recombination. Genome Biol Evol. 12(12):2508–2520.

Watterson GA. 1975. On the number of segregating sites in genetical models without recombination. Theor Popul Biol. 7(2):256–276.

Weir BS, Cockerham CC. 1984. Estimating F-statistics for the analysis of population structure. Evolution. 38(6):1358–1370.

Wenseleers T, Billen J. 2000. No evidence for *Wolbachia*-induced parthenogenesis in the social Hymenoptera. J Evol Biol. 13(2):277–280.

Werren JH. 1998. *Wolbachia* and speciation. In: Endless Forms: Species and Speciation. Oxford University Press. pp. 245–260.

Werren JH, Baldo L, Clark ME. 2008. *Wolbachia*: master manipulators of invertebrate biology. Nat Rev Microbiol. 6(10):741–751.

Williams GC. 1966. Natural selection, the costs of reproduction, and a refinement of Lack’s principle. Am Nat. 100(916):687–690.

Zhang YM, et al. 2020. UCE data reveal multiple origins of rose gallers in North America: globa phylogeny of *Diplolepis* Geoffroy (Hymenoptera: Cynipidae). Mol Phylogenet Evol. 153:106949.

