## Supplementary Material for "Population genomics of the mostly thelytokous *Diplolepis rosae* (Linnaeus, 1758) (Hymenoptera: Cynipidae) reveals population-specific selection for sex"

**SUPPORTING INFORMATION**

**Tables**

**Table S1.** Quality of the *Diplolepis rosae* genome assembly.

| Parameter | Value |
|---|---|
| # contigs (>= 0 bp) | 757 |
| # contigs (>= 1000 bp) | 729 |
| # contigs (>= 5000 bp) | 569 |
| # contigs (>= 10000 bp) | 429 |
| # contigs (>= 25000 bp) | 326 |
| # contigs (>= 50000 bp) | 285 |
| Total length (>= 0 bp) | 760724616 |
| Total length (>= 1000 bp) | 760705830 |
| Total length (>= 5000 bp) | 760203332 |
| Total length (>= 10000 bp) | 759172802 |
| Total length (>= 25000 bp) | 757619231 |
| Total length (>= 50000 bp) | 756165228 |
| Largest contig, bp | 33459371 |
| Total length, bp | 760724616 |
| % GC bases | 33.43 |
| N50 | 7663408 |
| N90 | 1376597 |
| L50 25 | 25 |
| L90 117 | 117 |
| # N's per 100 kbp | 0.00 |

**Table S2**. Classification and percentage of repetitive sequences in the *Diplolepis rosae* genome.

| Class of repetitive sequence | Number of elements † | Length occupied, bp | % of genome assembly |
|---|---|---|---|
| ***Retroelements:*** | 125770 | 86726618 | 11.40 |
| *SINEs:* | 10149 | 1528961 | 0.20 |
| Penelope | 39915 | 16183944 | 2.13 |
| *LINEs:* | 66655 | 38394411 | 5.05 |
| CRE/SLACS | 0 | 0 | 0 |
| L2/CR1/Rex | 0 | 0 | 0 |
| R1/LOA/Jockey | 21111 | 19220324 | 2.53 |
| R2/R4/NeSL | 872 | 334972 | 0.04 |
| RTE/Bov-B | 2151 | 558997 | 0.07 |
| L1/CIN4 | 0 | 0 | 0 |
| *LTR elements:* | 48966 | 46803246 | 6.15 |
| BEL/Pao | 2918 | 2652590 | 0.35 |
| Ty1/Copia | 2178 | 1798818 | 0.24 |
| Gypsy/DIRS1 | 43259 | 41907222 | 5.51 |
| Retroviral | 0 | 0 | 0 |
| ***DNA transposons:*** | 176423 | 65850608 | 8.66 |
| *hobo-Activator* | 13608 | 8583491 | 1.13 |
| *Tc1-IS630-Pogo* | 101005 | 38660213 | 5.08 |
| *En-Spm* | 0 | 0 | 0 |
| *MuDR-IS905* | 0 | 0 | 0 |
| *PiggyBac* | 20152 | 4871256 | 0.64 |
| *Tourist/Harbinger* | 1306 | 206185 | 0.03 |
| *Other (Mirage, P-element, Transib)* | 6541 | 2519567 | 0.33 |
| ***Rolling-circles*** | 1558 | 288437 | 0.04 |

| | | | |
|---|---|---|---|
| ***Unclassified*** | 1198129 | 368923706 | 48.50 |
| ***Small RNA*** | 7582 | 822276 | 0.11 |
| ***Satellites*** | 378 | 84139 | 0.01 |
| ***Simple repeats*** | 78424 | 3534451 | 0.46 |
| ***Low complexity*** | 15988 | 798635 | 0.10 |

† Most repeats fragmented by insertions or deletions have been counted as one element.

**Table S3**. Akaike Information Criterion (AIC) values for examined demographic scenarios for *Diplolepis rosae*.

| Model | AIC |
|---|---|
| bottlegrowth_split_no_migration | 74419.2 |
| bottlegrowth_split_migration | 79152.6 |
| split_no_migration | 144643.8 |
| split_asymmetric_migration | 145062.8 |
| split_symmetric_migration | 233881.6 |
| isolation_no_migration | 154600.6 |
| isol_asymmetric_migration | 146823.6 |
| isol_symmetric_migration | 159047.4 |
| exp_pop_growth_size_change_split_no_migration | 372553.8 |
| exp_pop_growth_size_change_split_asymmetric_migration | 371429.2 |
| exp_pop_growth_size_change_split_symmetric_migration | 371833.6 |

**Table S4**. Parameter estimations obtained by approximate Bayesian computation for the demographic model "Bottleneck_growth_split" in *Diplolepis rosae*.

| Parameter | N1 | N2 | proportion | Tbot | Tsplit | μ |
|---|---|---|---|---|---|---|
| Prediction error | 0.96 | 1.1 | 0.76 | 1.0 | 0.96 | 0.88 |
| Estimation, median | 511 | 494 | 0.62 | 1780 | 849 | 5.5e-7 |
| Units | Number of diploid individuals | | - | Generations ago | | per bp per generation |

Estimation is considered correct if its prediction value is less than 1. N1, N2: effective population sizes of population 1 and population 2, respectively; proportion: ratio between a number of individuals of *D. rosae* from a population after a bottleneck to an ancestral one; Tbot, Tsplit: bottleneck and split time, respectively; μ: mutation rate.

**Table S5.** Gene ontology terms showing outlier composite score values in the two *Diplolepis rosae* populations detected by Gene Set Enrichment Analysis.

| Composite score (CS) outliers in *D. rosae* lineages | Significant GO annotations |
|---|---|
| Negative (CS < -0.5) population 1 | GO:0046692: sperm competition (BP): 8/29 (p < 0.0001)<br>GO:0007320: insemination (BP): 8/32 (p < 0.0001)<br>GO:0007620: copulation (BP): 8/41 (p < 0.0001)<br>GO:0008237: metallopeptidase activity (MF): 11/123 (p = 0.000253)<br>GO:0004222: metalloendopeptidase activity (MF): 9/75 (p = 0.000253) |
| Negative (CS < -0.5) population 2 | GO:0071105: response to interleukin-1 (BP): 6/8 (p = 0.000278)<br>GO:0071348: cellular response to interleukin-11 (BP): 6/8 (p = 0.000278)<br>GO:1903544: response to butyrate (BP): 6/8 (p = 0.000278)<br>GO:1903545: cellular response to butyrate (BP): 6/8 (p = 0.000278)<br>GO:2001028: positive regulation of endothelial cell (BP): 7/13 (p = 0.000448)<br>GO:0099641: anterograde axonal protein transport (BP): 6/9 (p = 0.000462)<br>GO:1905907: negative regulation of amyloid fibril (BP): 6/9 (p = 0.000462)<br>GO:0032463: negative regulation of protein homooligomerization (BP): 7/14 (p = 0.000481)<br>GO:2001026: regulation of endothelial cell chemotaxis (BP): 7/14 (p = 0.000481)<br>GO:0038033: positive regulation of endothelial cell (BP): 6/10 (p = 0.000711)<br>GO:0042595: behavioral response to starvation (BP): 6/10 (p = 0.000711)<br>GO:1905906: regulation of amyloid fibril formation (BP): 7/16 (p = 0.00113)<br>GO:0032460: negative regulation of protein oligomer (BP): 7/17 (p = 0.00139)<br>GO:0032462: regulation of protein homooligomerization (BP): 7/17 (p = 0.00139)<br>GO:0035767: endothelial cell chemotaxis (BP): 7/17 (p = 0.00139)<br>GO:0042026: protein refolding (BP): 7/17 (p = 0.00139)<br>GO:0038089: positive regulation of cell migration (BP): 6/12 (p = 0.00190)<br>GO:0032459: regulation of protein oligomerization (BP): 8/25 (p = 0.00202)<br>GO:1905383: protein localization to presynapse (BP): 6/13 (p = 0.00305)<br>GO:1990000: amyloid fibril formation (BP): 7/20 (p = 0.00401)<br>GO:0007021: tubulin complex assembly (BP): 6/14 (p = 0.00468)<br>GO:0032731: positive regulation of interleukin-1 (BP): 6/15 (p = 0.00718)<br>GO:1902176: negative regulation of oxidative stress (BP): 7/22 (p = 0.00718)<br>GO:0051016: barbed-end actin filament capping (BP): 6/16 (p = 0.00909)<br>GO:0098840: protein transport along microtubule (BP): 6/16 (p = 0.00909)<br>GO:0099118: microtubule-based protein transport (BP): 6/16 (p = 0.00909)<br>GO:0099640: axo-dendritic protein transport (BP): 6/16 (p = 0.00909)<br>GO:1990776: response to angiotensin (BP): 8/32 (p = 0.0100)<br>GO:0009631: cold acclimation (BP): 6/17 (p = 0.0122)<br>GO:0032732: positive regulation of interleukin-1 (BP): 6/17 (p = 0.0122) |

| | |
|---|---|
| | GO:0071679: commissural neuron axon guidance (BP): 6/18 ($p = 0.0169$)<br>GO:0032651: regulation of interleukin-1 beta product (BP): 7/26 ($p = 0.0169$)<br>GO:1902175: regulation of oxidative stress-induced (BP): 7/26 ($p = 0.0169$)<br>GO:0032611: interleukin-1 beta production (BP): 7/28 ($p = 0.0277$)<br>GO:0002931: response to ischemia (BP): 7/29 ($p = 0.0333$)<br>GO:0032652: regulation of interleukin-1 production (BP): 7/29 ($p = 0.0333$)<br>GO:0035994: response to muscle stretch (BP): 6/22 ($p = 0.0450$)<br>GO:0051693: actin filament capping (BP): 6/22 ($p = 0.0450$)<br>GO:0008089: anterograde axonal transport (BP): 8/41 ($p = 0.0450$)<br>GO:1900408: negative regulation of cellular response (BP): 8/41 ($p = 0.0450$)<br>GO:1903202: negative regulation of oxidative stress (BP): 8/41 ($p = 0.0450$)<br>GO:0032612: interleukin-1 production (BP): 7/31 ($p = 0.0450$)<br>GO:0008426: protein kinase C inhibitor activity (MF): 7/10 ($p = 0.000159$)<br>GO:0005212: structural constituent of eye lens (MF): 6/11 ($p = 0.00776$)<br>GO:0097512: cardiac myofibril (CC): 6/10 ($p = 0.00527$) |
| Positive (CS > 0.5) population 1 | GO:0071679: commissural neuron axon guidance (BP): 6/18 ($p = 0.0139$) |
| Positive (CS > 0.5) population 2 | GO:0046692: sperm competition (BP): 9/29 ($p < 0.0001$)<br>GO:0007320: insemination (BP): 9/32 ($p < 0.0001$)<br>GO:0007620: copulation (BP): 9/41 ($p < 0.0001$)<br>GO:0007617: mating behavior (BP): 10/167 ($p = 0.00940$)<br>GO:0044706: multi-multicellular process (BP): 9/152 ($p = 0.0271$)<br>GO:0007618: mating (BP): 9/160 ($p = 0.0341$)<br>GO:0008237: metalloendopeptidase activity (MF): 75/10 ($p < 0.0001$)<br>GO:0004222: metallopeptidase activity (MF): 11/123 ($p < 0.0001$)<br>GO:0004175: endopeptidase activity (MF): 11/203 ($p = 0.00780$)<br>GO:0004175: peptidase activity (MF): 13/340 ($p = 0.0300$) |

Results are presented as “Gene ontology (GO) identifier: annotation (GO): number of detected significant genes / total number of genes in a full gene set (adjusted p-value, Fisher’s exact test)”. BP: ‘Biological Process’ ontology. MF: ‘Molecular Function’ ontology. CC: ‘Cellular component’ ontology.

**Table S6.** Coverage *Wolbachia* supergroup A and B bins (contigs built after metagenome assembly and binning) recovered from *D. rosae* Illumina reads.

| Population | *D. rosae* individual | bin | Coverage | Identified *Wolbachia* Supergroup (A or B) | *D. rosae* genome coverage | Normalized *Wolbachia* coverage |
|---|---|---|---|---|---|---|
| 1 | 078 | bin.1.fa | 9.82822 | | 44.8897 | |
| 1 | 078 | bin.2.fa | 36.2733 | | | |
| 1 | 078 | bin.3.fa | 17.3943 | | | |
| 1 | 078 | bin.4.fa | 5.21198 | | | |
| 1 | 078 | bin.5.fa | 13.0497 | A and B | | 0.29 |
| 1 | 082 | bin.1.fa | 9.17111 | | 58.1444 | |
| 1 | 082 | bin.2.fa | 14.354 | B | | 0.25 |
| 1 | 082 | bin.3.fa | 9.34931 | | | |
| 1 | 082 | bin.4.fa | 7.32544 | | | |
| 1 | 082 | bin.5.fa | 13.8626 | | | |
| 1 | 082 | bin.6.fa | 14.8484 | | | |
| 1 | 082 | bin.7.fa | 28.1723 | | | |
| 1 | 082 | bin.8.fa | 9.44117 | | | |
| 2 | 117 | bin.1.fa | 9.9021 | | 25.9941 | |
| 2 | 117 | bin.2.fa | 23.8607 | A and B | | 0.92 |
| 2 | 117 | bin.3.fa | 18.5399 | | | |
| 2 | 117 | bin.4.fa | 43.5673 | | | |
| 1 | 126 | bin.1.fa | 24.5021 | | 27.1813 | |
| 1 | 126 | bin.2.fa | 4.5765 | | | |
| 1 | 126 | bin.3.fa | 13.4358 | | | |
| 1 | 126 | bin.4.fa | 6.61558 | | | |
| 1 | 126 | bin.5.fa | 28.1247 | | | |
| 1 | 126 | bin.6.fa | 13.6376 | B | | 0.50 |
| 1 | 126 | bin.7.fa | 26.043 | A | | 0.96 |
| 1 | 219 | bin.1.fa | 46.5367 | | 119.241 | |
| 1 | 219 | bin.2.fa | 23.4625 | | | |
| 1 | 219 | bin.3.fa | 10.7127 | | | |
| 1 | 219 | bin.4.fa | 42.6195 | | | |
| 1 | 219 | bin.5.fa | 107.281 | B | | 0.90 |
| 1 | 219 | bin.6.fa | 17.9632 | | | |
| 2 | 288 | bin.1.fa | 8.33563 | | 38.8936 | |
| 2 | 288 | bin.2.fa | 8.5904 | | | |
| 2 | 288 | bin.3.fa | 93.9574 | A | | 2.4 |

| | | | | | | |
|---|---|---|---|---|---|---|
| 2 | 288 | bin.4.fa | 122.614 | B | | 3.2 |
| 2 | 288 | bin.5.fa | 148.543 | B | | 3.8 |
| 1 | 312 | bin.1.fa | 251.813 | B | 34.1124 | 7.4 |
| 1 | 312 | bin.2.fa | 8.66442 | | | |
| 2 | 330 | bin.1.fa | 7.54022 | | 36.7328 | |
| 2 | 330 | bin.2.fa | 7.33231 | | | |
| 2 | 330 | bin.3.fa | 8.48644 | | | |
| 1 | 464 | bin.1.fa | 5.54094 | | 29.4562 | |
| 1 | 464 | bin.2.fa | 348.081 | | | |
| 1 | 464 | bin.3.fa | 14.0629 | | | |
| 1 | 464 | bin.4.fa | 308.676 | B | | 10.5 |
| 1 | 464 | bin.5.fa | 12.6503 | | | |
| 1 | 464 | bin.6.fa | 13.2359 | | | |
| 1 | 576 | bin.10.fa | 84.098 | B | 38.3787 | 2.2 |
| 1 | 576 | bin.11.fa | 16.5527 | B | | 0.43 |
| 1 | 576 | bin.12.fa | 11.6308 | | | |
| 1 | 576 | bin.13.fa | 11.2718 | | | |
| 1 | 576 | bin.1.fa | 12.8816 | | | |
| 1 | 576 | bin.2.fa | 11.2316 | | | |
| 1 | 576 | bin.3.fa | 11.9606 | | | |
| 1 | 576 | bin.4.fa | 11.3751 | | | |
| 1 | 576 | bin.5.fa | 11.9592 | | | |
| 1 | 576 | bin.6.fa | 7.63389 | | | |
| 1 | 576 | bin.7.fa | 7.69986 | | | |
| 1 | 576 | bin.8.fa | 11.5755 | | | |
| 1 | 576 | bin.9.fa | 11.18 | | | |
| 2 | 580 | bin.1.fa | 9.88938 | | 39.3057 | |
| 2 | 580 | bin.2.fa | 13.0791 | | | |
| 2 | 580 | bin.3.fa | 10.38 | | | |
| 2 | 580 | bin.4.fa | 8.82744 | | | |
| 2 | 580 | bin.5.fa | 10.5102 | | | |
| 2 | 580 | bin.6.fa | 7.80195 | | | |
| 2 | 580 | bin.7.fa | 12.2507 | | | |
| 2 | 580 | bin.8.fa | 152.488 | B | | 3.9 |
| 1 | 608 | bin.1.fa | 89.8094 | B | 36.2919 | 2.5 |
| 1 | 608 | bin.2.fa | 20.7932 | | | |
| 1 | 608 | bin.3.fa | 8.68584 | | | |

| | | | | | | |
|---|---|---|---|---|---|---|
| 1 | 608 | bin.4.fa | 190.495 | A | | 5.2 |
| 1 | 608 | bin.5.fa | 8.2291 | | | |
| 2 | 623 | bin.1.fa | 474.038 | B | 33.7402 | 14.0 |
| 2 | 623 | bin.2.fa | 554.346 | | | |
| 2 | 623 | bin.3.fa | 6.25831 | | | |
| 1 | 630 | bin.1.fa | 19.6523 | | 32.8526 | |
| 1 | 630 | bin.2.fa | 16.5952 | | | |
| 1 | 630 | bin.3.fa | 106.839 | B | | 0.60 |
| 1 | 630 | bin.4.fa | 33.1374 | | | |
| 1 | 630 | bin.5.fa | 7.15868 | | | |
| 1 | 630 | bin.6.fa | 7.00814 | | | |
| 2 | 652 | bin.1.fa | 146.607 | B | 46.3449 | 3.2 |
| 2 | 652 | bin.2.fa | 8.9036 | | | |
| 2 | 652 | bin.3.fa | 8.3211 | | | |

**Table S6.** Coverage of *Wolbachia* supergroup A and B contigs built after metagenome assembly and binning (bins) recovered from *Diplolepis rosae* Illumina reads.

| Species | Sample name | Stage | Number of individuals per sample | Tissue | Sex | Host plant | Isolation source | Collection date | Latitude | Longitude |
|---|---|---|---|---|---|---|---|---|---|---|
| *Diplolepis rosae* | ESE-078 | Imago | 6 | Whole individual | Female | *Rosa canina* | Gall | 2020-05-12 | 48.701996 | 2.173458 |
| *Diplolepis rosae* | ESE-082 | Imago | 7 | Whole individual | Female | *Rosa canina* | Gall | 2020-05-13 | 48.703719 | 2.083596 |
| *Diplolepis rosae* | ESE-117 | Imago | 8 | Whole individual | Female | *Rosa canina* | Gall | 2020-04-04 | 45.758762 | 6.159383 |
| *Diplolepis rosae* | ESE-126 | Imago | 8 | Whole individual | Female | *Rosa canina* | Gall | 2020-06-14 | 47.249305 | 6.073626 |
| *Diplolepis rosae* | ESE-219 | Imago (Illumina) Larva (Nanopore) | 30 | Whole individual | Female | *Rosa canina* | Gall | 2020-08-10 | 46.422178 | 1.353230 |
| *Diplolepis rosae* | ESE-288 | Larva | 7 | Whole individual | Female | *Rosa canina* | Gall | 2020-08-26 | 47.324787 | 3.792692 |
| *Diplolepis rosae* | ESE-312 | Larva | 10 | Whole individual | Female | *Rosa canina* | Gall | 2020-09-02 | 50.006474 | 4.743896 |
| *Diplolepis rosae* | ESE-330 | Larva | 20 | Whole individual | Female | *Rosa canina* | Gall | 2020-09-05 | 44.527561 | 5.016034 |
| *Diplolepis rosae* | ESE-349 | Larva | 10 | Whole individual | Female | *Rosa canina* | Gall | 2020-09-08 | 45.620130 | 3.057751 |
| *Diplolepis rosae* | ESE-464 | Larva | 10 | Whole individual | Female | *Rosa canina* | Gall | 2020-09-23 | 43.447913 | 6.239238 |
| *Diplolepis rosae* | ESE-559 | Larva | 12 | Whole individual | Female | *Rosa canina* | Gall | 2020-09-10 | 43.967949 | 3.405437 |
| *Diplolepis rosae* | ESE-576 | Larva | 10 | Whole individual | Female | *Rosa canina* | Gall | 2020-10-12 | 43.033091 | 1.003795 |
| *Diplolepis rosae* | ESE-580 | Larva | 10 | Whole individual | Female | *Rosa canina* | Gall | 2020-10-09 | 42.702320 | 2.564203 |
| *Diplolepis rosae* | ESE-608 | Larva | 8 | Whole individual | Female | *Rosa canina* | Gall | 2020-10-20 | 44.589800 | -0.454849 |
| *Diplolepis rosae* | ESE-623 | Larva | 7 | Whole individual | Female | *Rosa canina* | Gall | 2020-10-20 | 43.655179 | -1.180453 |
| *Diplolepis rosae* | ESE-630 | Larva | 10 | Whole individual | Female | *Rosa canina* | Gall | 2020-10-20 | 45.150354 | 0.529976 |
| *Diplolepis rosae* | ESE-652 | Larva | 5 | Whole individual | Female | *Rosa canina* | Gall | 2020-12-27 | 47.309109 | -3.067183 |

### Figures

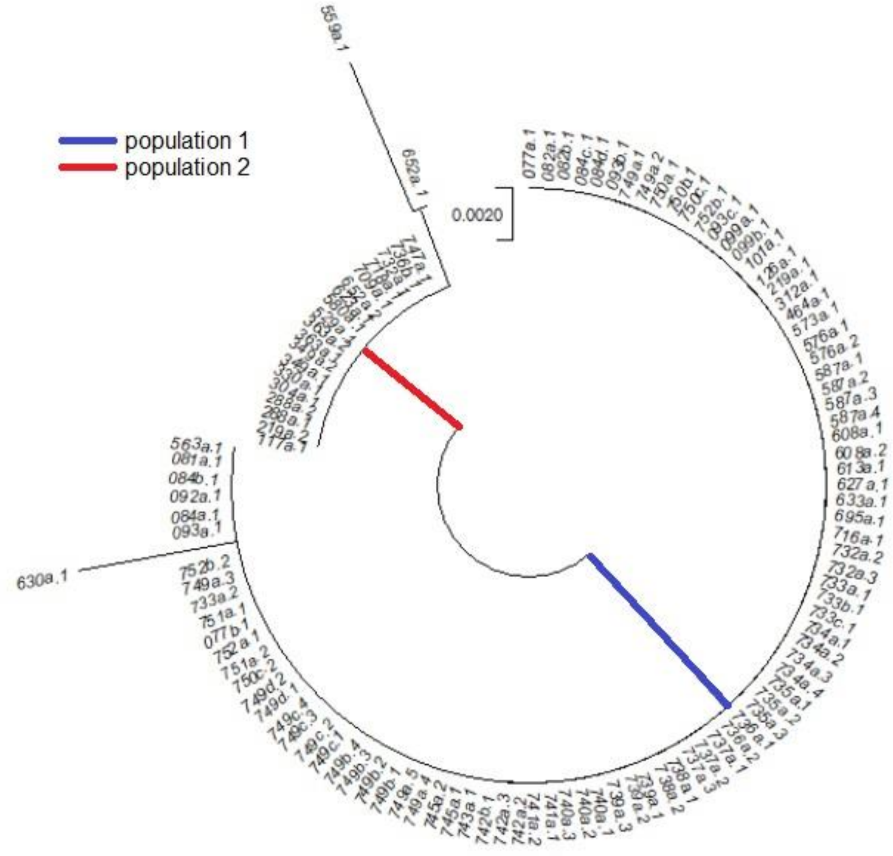

**Fig. S1**. Phyloogram of *Diplolepis rosae* genotypes collected in France based on one marker. Two branches correspond to the two populations.

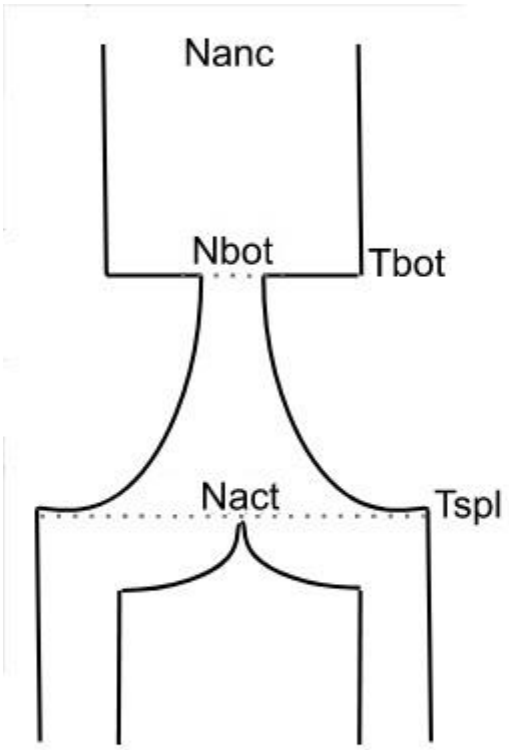

**Fig S2**. Schema describing the best-fitted model for the demographic scenario of *Diplolepis rosae*: a bottleneck of an ancestral population followed by exponential growth then split into two populations with no gene flow between them. Nanc, Nbot, Nact: population sizes (ancestral, after the bottleneck, just before the split, respectively); Tbot, Tspl: times (bottleneck, split, respectively). Line sizes are scaled relatively to *dadi* estimates: Nbot/Nanc = 0.13; Nact/Nanc = 2.4; bottleneck time = 0.22; split time = 0.20; time unit is 2*Neff generations ago; Neff: effective population size (diploid individuals).

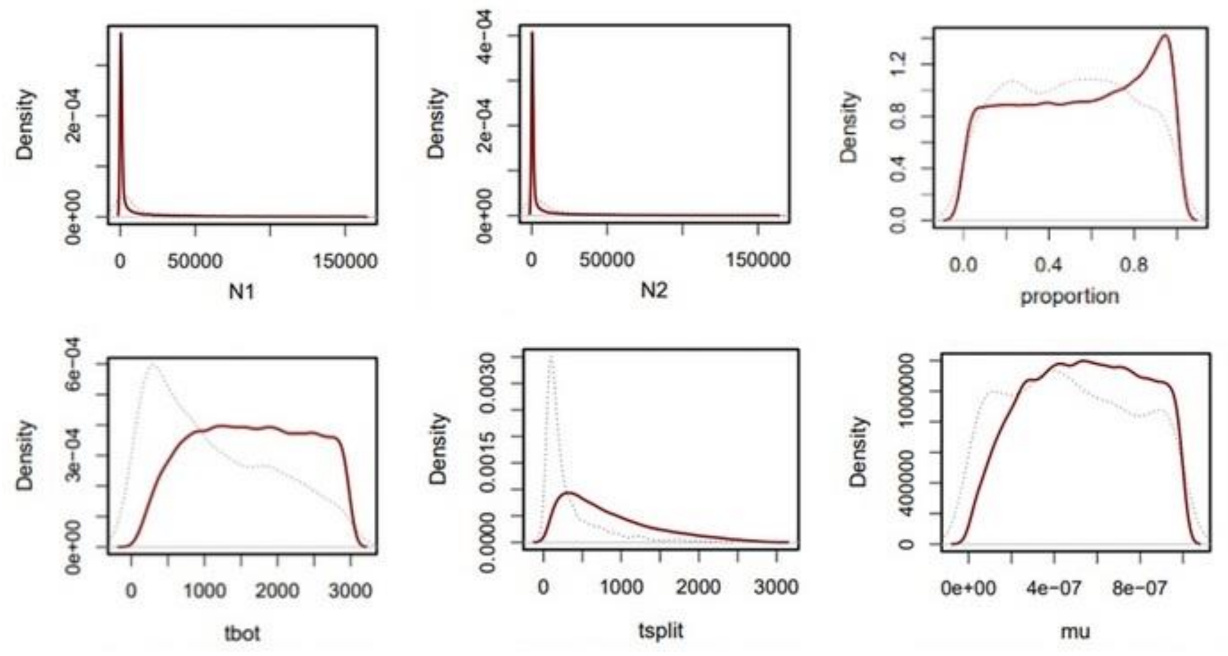

**Fig. 3**. Parameter inference of the simulated demographic model "Bottleneck_growth_split" in *Diplolepis rosae* using approximate Bayesian computation (ABC). Grey line: given distribution of simulated model parameters (priors). Red line: distribution-accepted model parameters (posteriors). N1, N2: effective population sizes of the first and the second populations, respectively; proportion: ratio between the number of individuals of *D. rosae* from a population after a bottleneck to an ancestral one; tbot, tsplit: bottleneck and split time, respectively; mu: mutation rate.

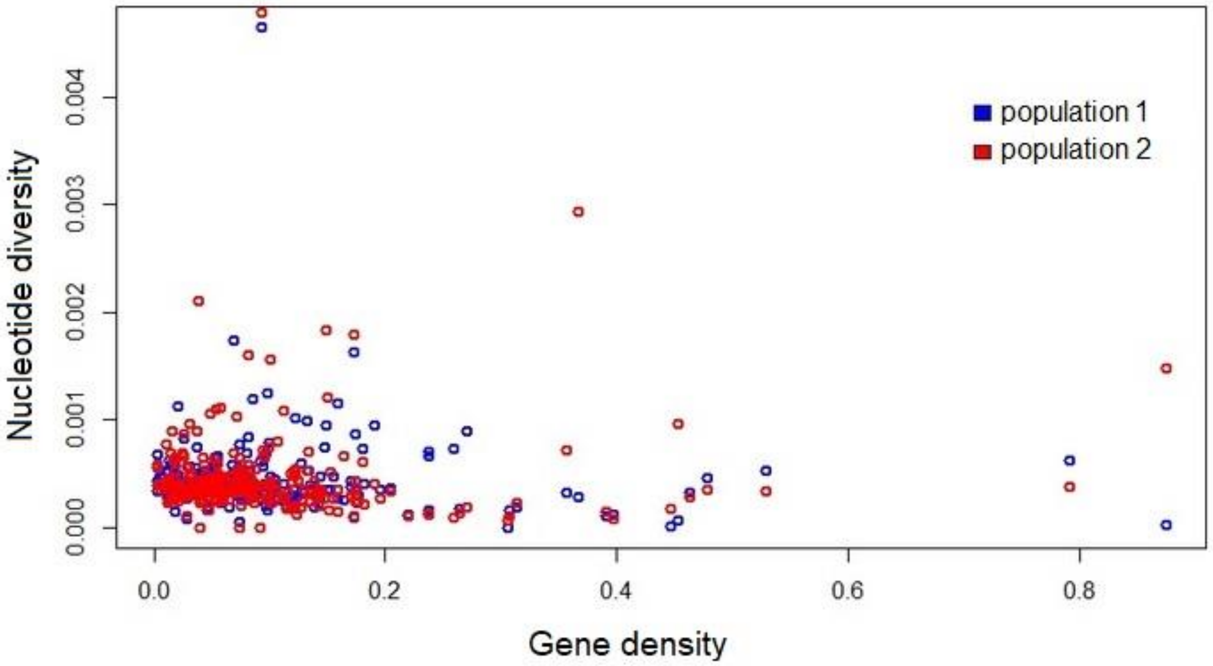

**Fig. S4**. Relationship between gene density (the proportion of nucleotides assigned to a protein coding sequence in a genome fragment) and nucleotide diversity ($\pi$) in *Diplolepis rosae*.

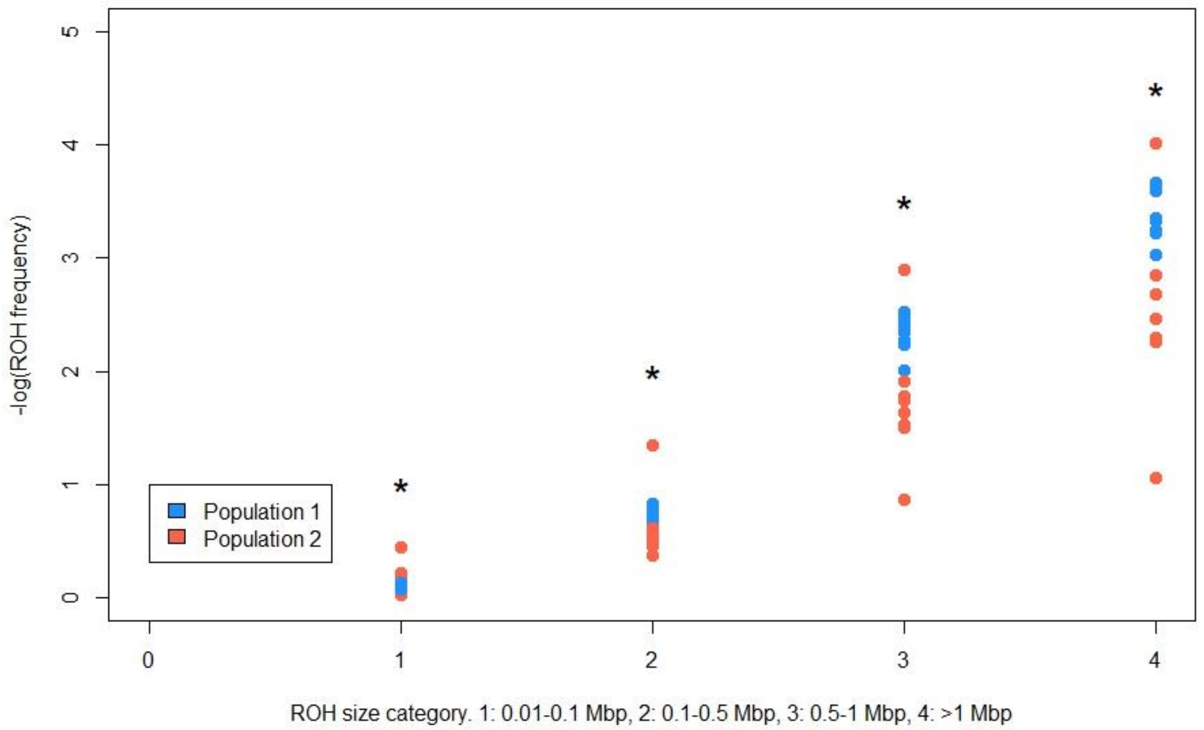

**Fig S5**. Frequency of runs of homozygosity (ROHs) of different size categories in the *Diplolepis rosae* genome. Each point represents one *D. rosae* individual. The ROH frequency is presented as the ratio between the number of ROHs from each size category and the total number of ROHs on a negative logarithmic scale. The asterisks indicate a significant difference in ROH frequencies between population 1 and population 2 (Mann–Whitney U test, $p < 0.05$). In the ROH size categories '0.5–1 Mb' and '> 1 Mb', the lowest values of −log(ROH frequency) correspond to the admixed individual *D. rosae-652* (**Fig. 1**).

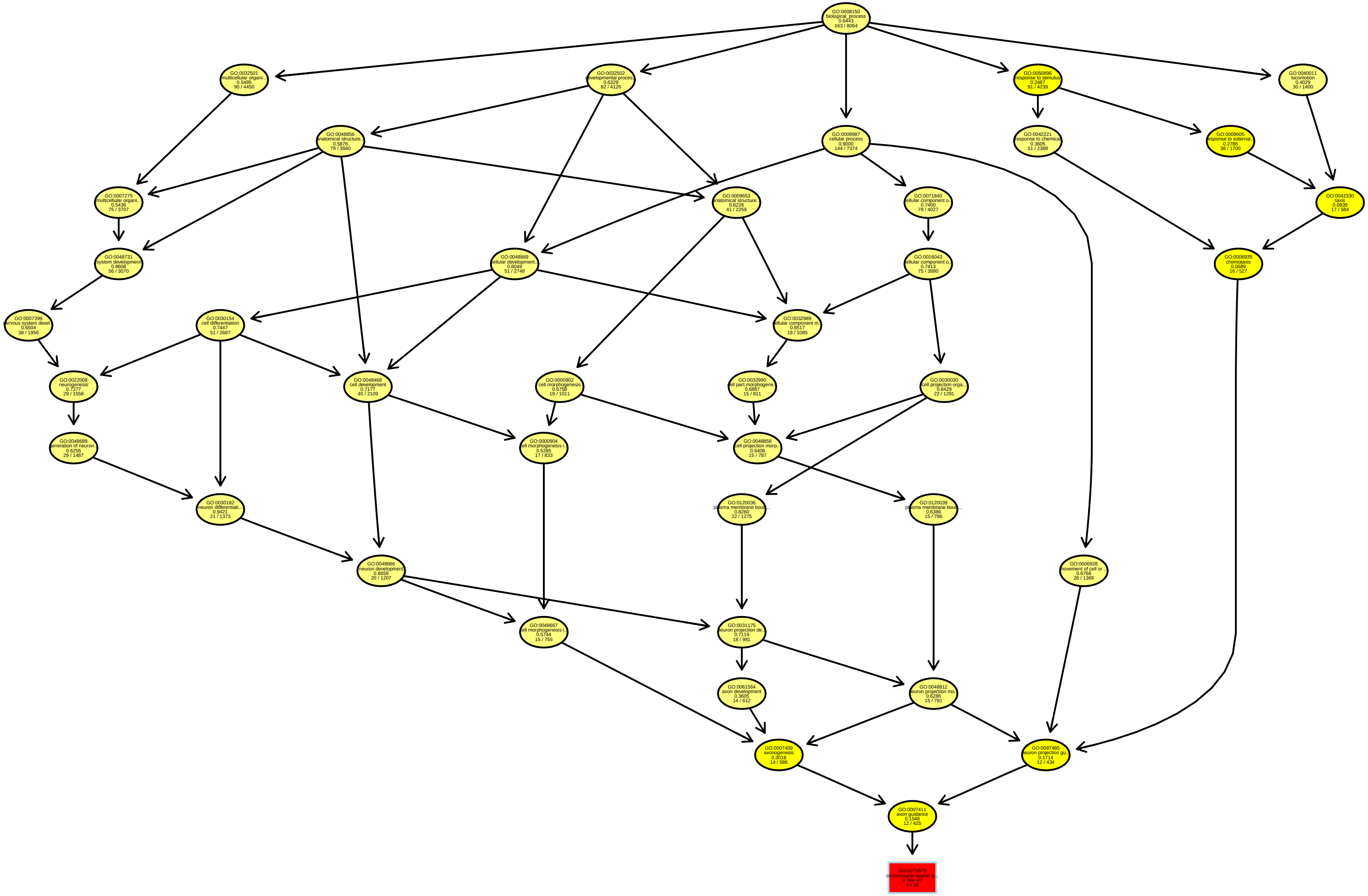

GO:0008150 biological_process 0.6443 163 / 8064
GO:0032501 multicellular organi... 0.5495 90 / 4450
GO:0032502 developmental proces... 0.6329 82 / 4126
GO:0009987 cellular process 0.9000 144 / 7374
GO:0050896 response to stimulus 0.2487 91 / 4239
GO:0040011 locomotion 0.4029 30 / 1400
GO:0048856 anatomical structure... 0.5876 79 / 3940
GO:0042221 response to chemical 0.3605 51 / 2388
GO:0009605 esponse to external... 0.2786 38 / 1700
GO:0007275 multicellular organi... 0.5436 75 / 3707
GO:0009653 anatomical structure... 0.8226 41 / 2259
GO:0071840 ellular component o... 0.7400 78 / 4027
GO:0042330 taxis 0.0839 17 / 584
GO:0048731 system development 0.8608 56 / 3070
GO:0048869 ellular development... 0.8049 51 / 2748
GO:0016043 ellular component o... 0.7413 75 / 3880
GO:0006935 chemotaxis 0.0689 16 / 527
GO:0007399 ervous system devel... 0.6504 38 / 1956
GO:0030154 cell differentiation 0.7447 51 / 2687
GO:0032989 ellular component m... 0.8517 18 / 1085
GO:0022008 neurogenesis 0.7277 29 / 1558
GO:0048468 cell development 0.7177 40 / 2109
GO:0000902 cell morphogenesis 0.6758 19 / 1011
GO:0032990 ell part morphogene... 0.6867 15 / 811
GO:0030030 cell projection orga... 0.8429 22 / 1291
GO:0048699 eneration of neuron... 0.6255 29 / 1487
GO:0000904 ell morphogenesis i... 0.5285 17 / 833
GO:0048858 cell projection morp... 0.6406 15 / 787
GO:0030182 neuron differentiati... 0.9421 21 / 1373
GO:0120036 plasma membrane boun... 0.8260 22 / 1275
GO:0120039 plasma membrane boun... 0.6386 15 / 786
GO:0048666 neuron development 0.8656 20 / 1207
GO:0006928 movement of cell or ... 0.6768 26 / 1369
GO:0048667 ell morphogenesis i... 0.5744 15 / 755
GO:0031175 euron projection de... 0.7119 18 / 981
GO:0061564 axon development 0.3605 14 / 612
GO:0048812 euron projection mo... 0.6286 15 / 781
GO:0007409 axonogenesis 0.3018 14 / 586
GO:0097485 euron projection gu... 0.1714 12 / 434
GO:0007411 axon guidance 0.1548 12 / 425
GO:0071679 commissural neuron a... 9.74e−07 6 / 18

**Fig S6.** Relationship in the 'Biological process' ontology between gene terms showing a positive outlier composite score in *Diplolepis rosae* population 1. Results are presented as Gene Ontology (GO) term, annotation, raw p-value (Fisher's exact test), number of detected significant genes in *D. rosae* / total number of genes in the full gene set.

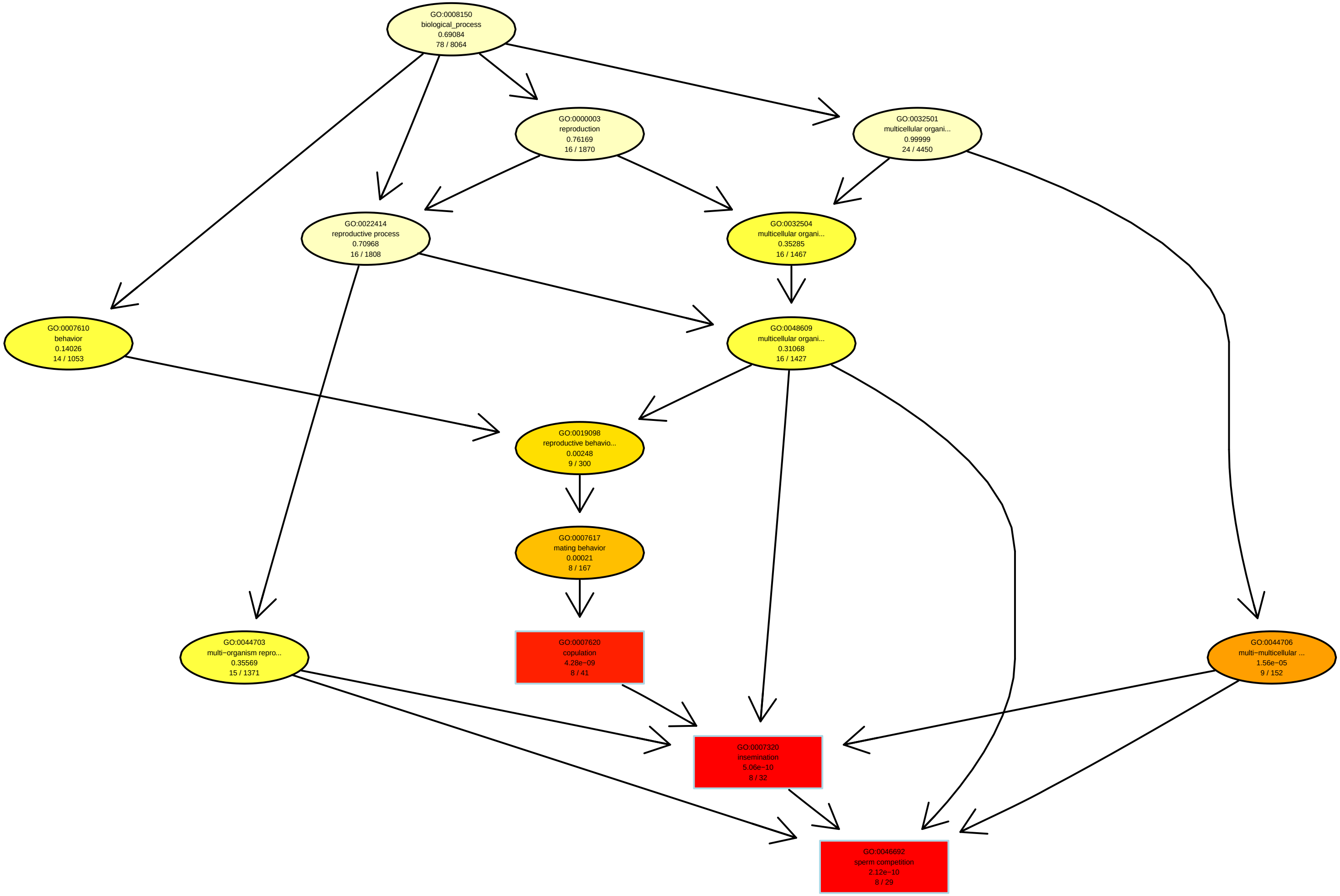
GO:0008150
biological_process
0.69084
78 / 8064
GO:0000003
reproduction
0.76169
16 / 1870
GO:0032501
multicellular organi...
0.99999
24 / 4450
GO:0022414
reproductive process
0.70968
16 / 1808
GO:0032504
multicellular organi...
0.35285
16 / 1467
GO:0007610
behavior
0.14026
14 / 1053
GO:0048609
multicellular organi...
0.31068
16 / 1427
GO:0019098
reproductive behavio...
0.00248
9 / 300
GO:0007617
mating behavior
0.00021
8 / 167
GO:0044703
multi−organism repro...
0.35569
15 / 1371
GO:0007620
copulation
4.28e−09
8 / 41
GO:0044706
multi−multicellular ...
1.56e−05
9 / 152
GO:0007320
insemination
5.06e−10
8 / 32
GO:0046692
sperm competition
2.12e−10
8 / 29

**Fig S7.** Relationship in the 'Biological process' ontology between gene terms showing negative outlier composite score in *Diplolepis rosae* population 1. Results are presented as Gene Ontology (GO) term, annotation, raw p-value (Fisher's exact test), number of detected genes in *D. rosae* / total number of genes in the full gene set.

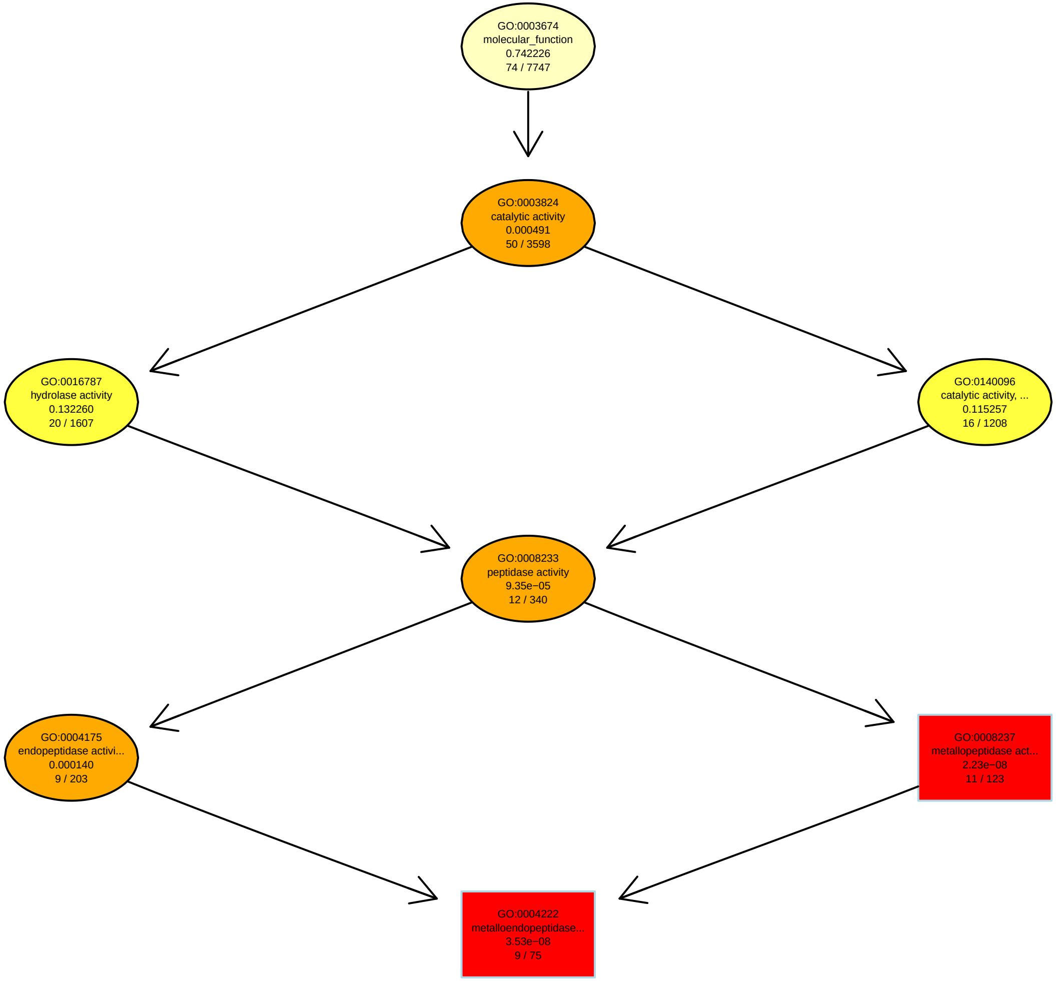
GO:0003674
molecular_function
0.742226
74 / 7747
GO:0003824
catalytic activity
0.000491
50 / 3598
GO:0016787
hydrolase activity
0.132260
20 / 1607
GO:0140096
catalytic activity, ...
0.115257
16 / 1208
GO:0008233
peptidase activity
9.35e−05
12 / 340
GO:0004175
endopeptidase activi...
0.000140
9 / 203
GO:0008237
metallopeptidase act...
2.23e−08
11 / 123
GO:0004222
metalloendopeptidase...
3.53e−08
9 / 75

**Fig S8.** Relationship in the 'Molecular Function' ontology between gene terms showing a negative outlier composite score in *Diplolepis rosae* population 1. Results are presented as Gene Ontology (GO) term, annotation, raw p-value (Fisher's exact test), number of detected genes in *D. rosae* / total number of genes in the full gene set.

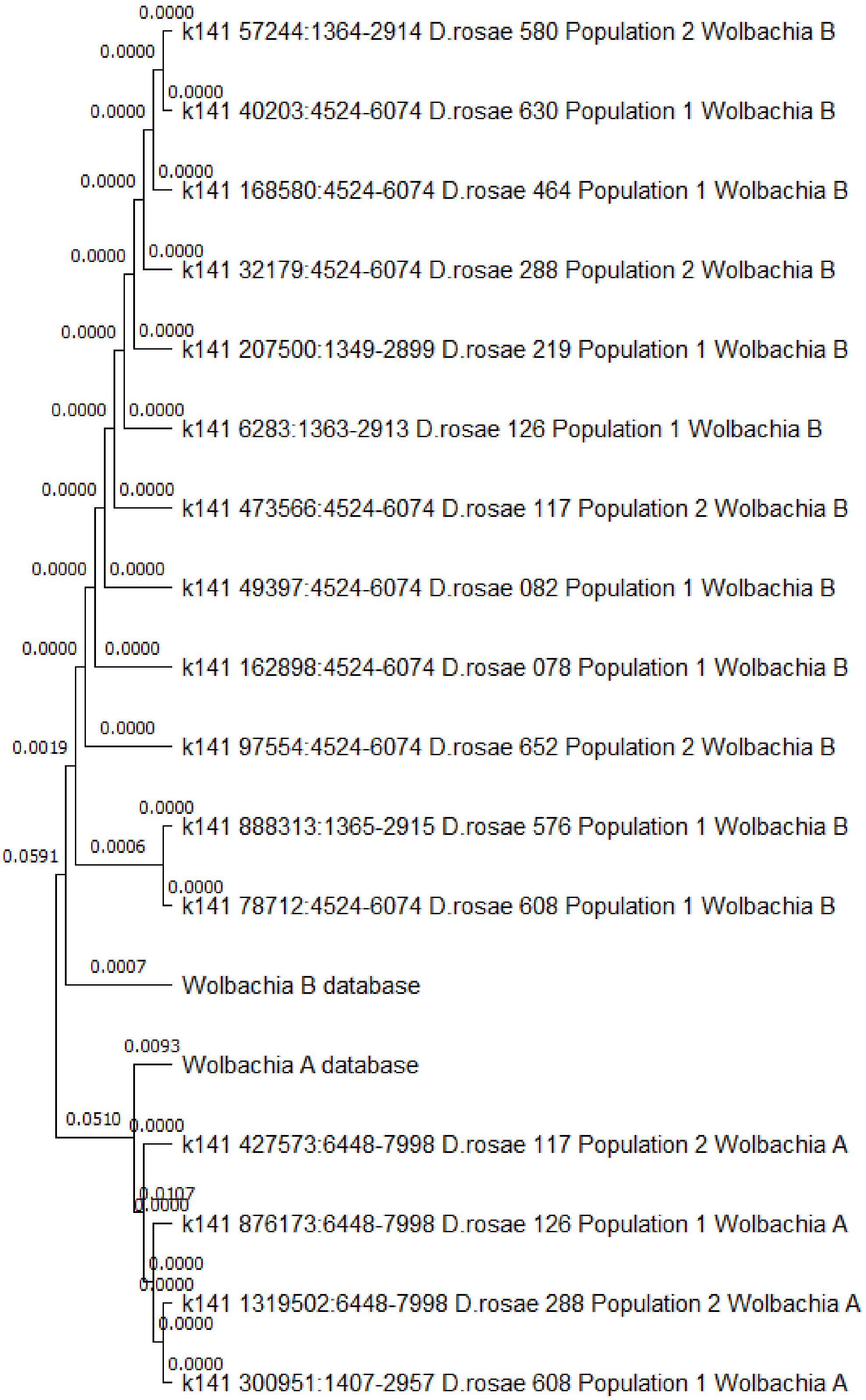

**Fig. S9**. Phyloogram of *Wolbachia* genotypes based on the *coxA* gene (Wang et al. 2020) recovered from *Diplolepis rosae* Illumina sequences. Each sample corresponds to *Wolbachia* contig identifier: start–end position of the gene_corresponding *D.rosae* sample name_*D. rosae* population_*Wolbachia* supergroup. The phylogram was constructed using the Maximum likelihood statistical method (Tamura–Nei substitution model (default)) by MEGA 11 (Tamura et al. 2021).

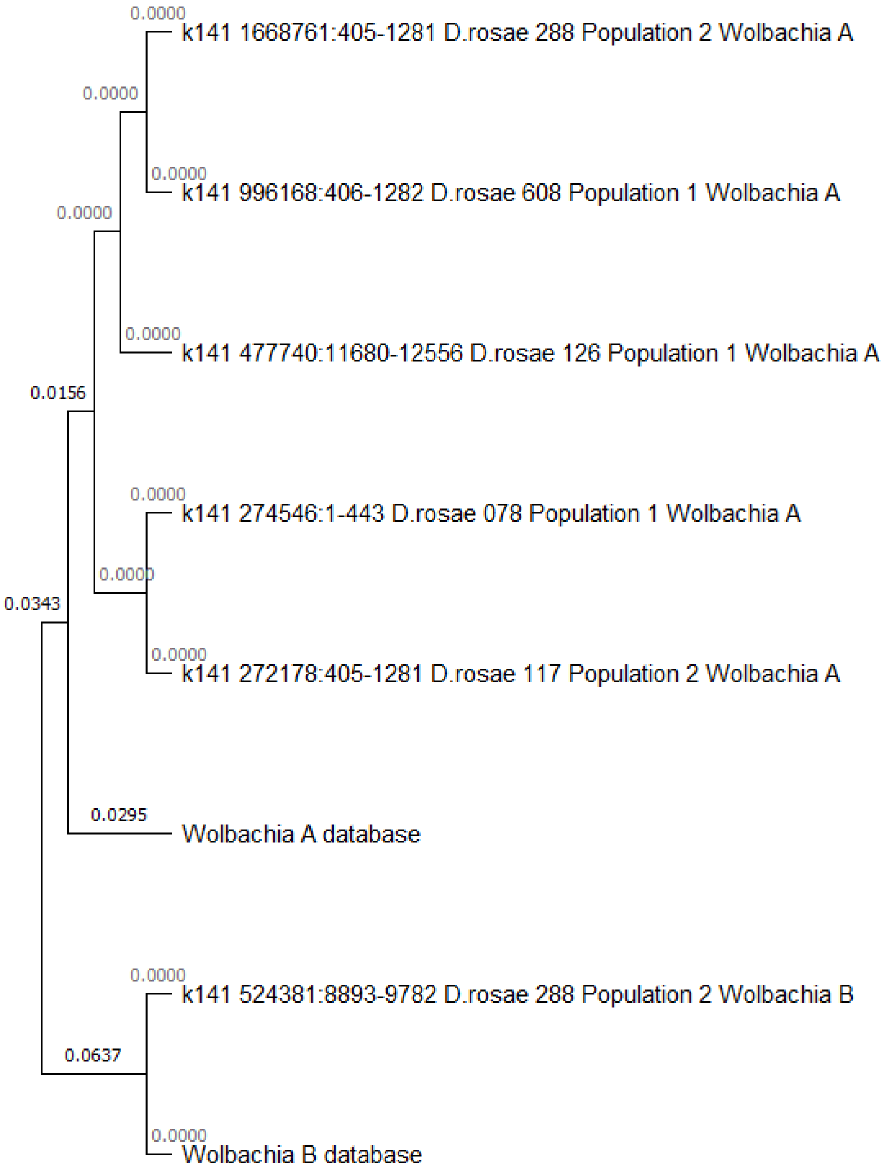

**Fig. S10**. Phyloogram of *Wolbachia* genotypes (*fbpA* gene, see details in Wang et al. 2020) recovered from *D. rosae* Illumina sequences. Each sample corresponds to *Wolbachia* contig identifier: start – end position of the gene_corresponding *D.rosae* sample name_*D. rosae* population_*Wolbachia* supergroup. The phylogram was constructed using the Maximum likelihood statistical method (Tamura-Nei substitution model (default)) by MEGA 11 (Tamura et al. 2021).

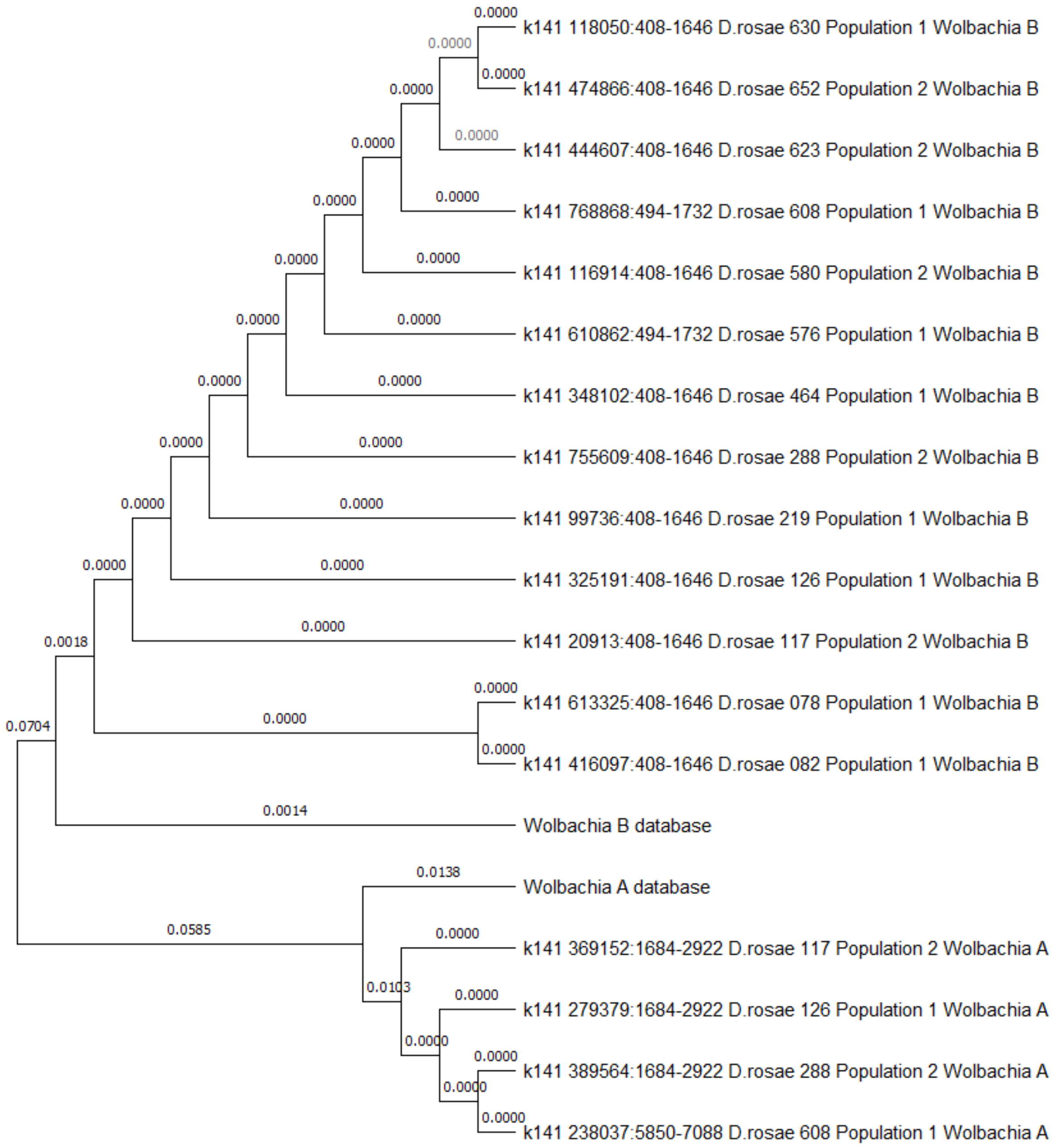

**Fig. S11**. Phyloogram of *Wolbachia* genotypes based on the *ftsZ* gene (Wang et al. 2020) recovered from *Diplolepis rosae* Illumina sequences. Each sample corresponds to *Wolbachia* contig identifier: start–end position of the gene_corresponding *D.rosae* sample name_*D. rosae* population_*Wolbachia* supergroup. The phylogram was constructed using the Maximum likelihood statistical method (Tamura–Nei substitution model (default)) by MEGA 11 (Tamura et al. 2021).

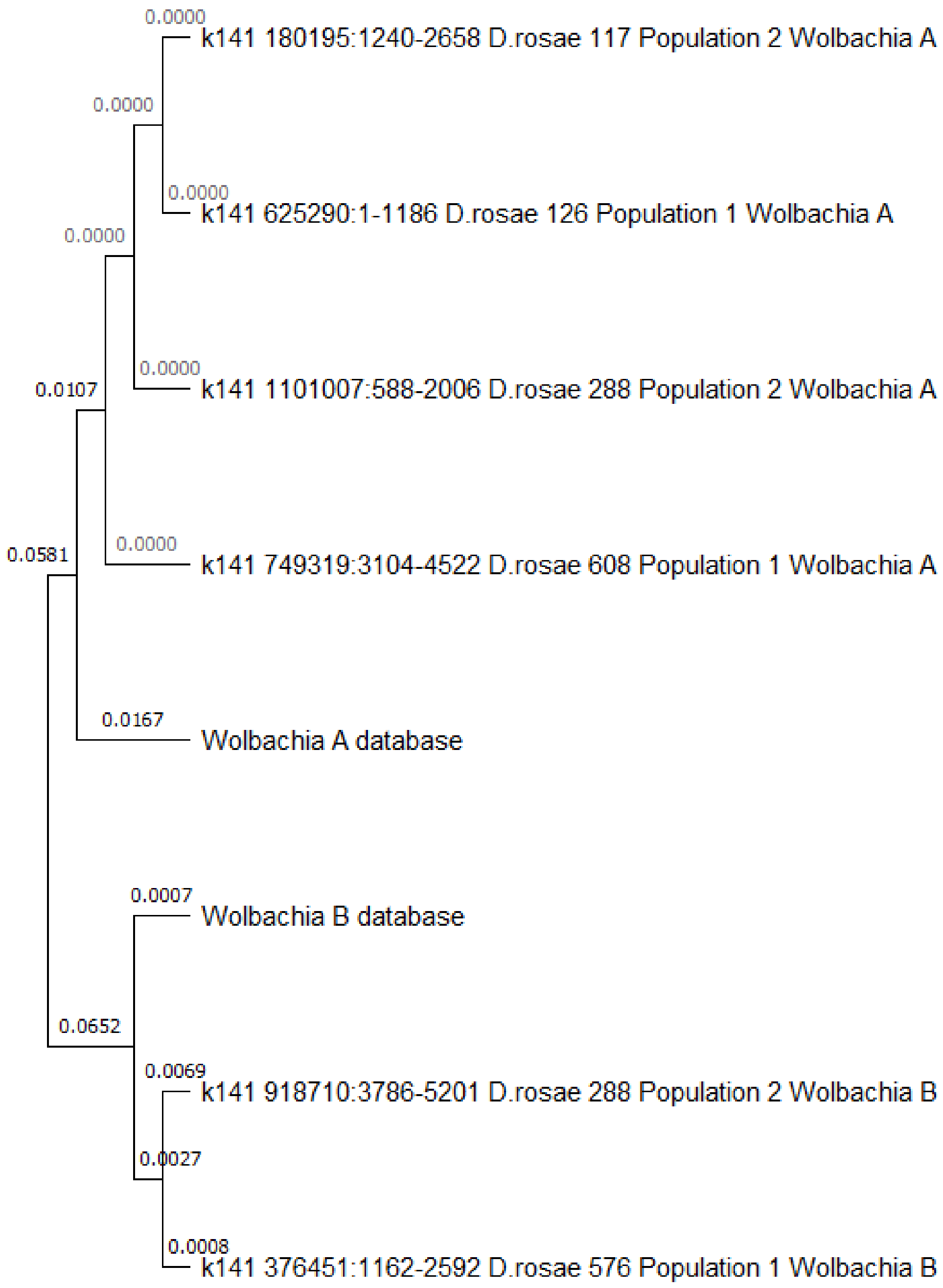

**Fig. S12**. Phyloogram of *Wolbachia* genotypes based on the *gatB* gene (Wang et al. 2020) recovered from *Diplolepis rosae* Illumina sequences. Each sample corresponds to *Wolbachia* contig identifier: start–end position of the gene corresponding *D.rosae* sample name_*D. rosae* population_*Wolbachia* supergroup. The phylogram was constructed using the Maximum likelihood statistical method (Tamura–Nei substitution model (default)) by MEGA 11 (Tamura et al. 2021).

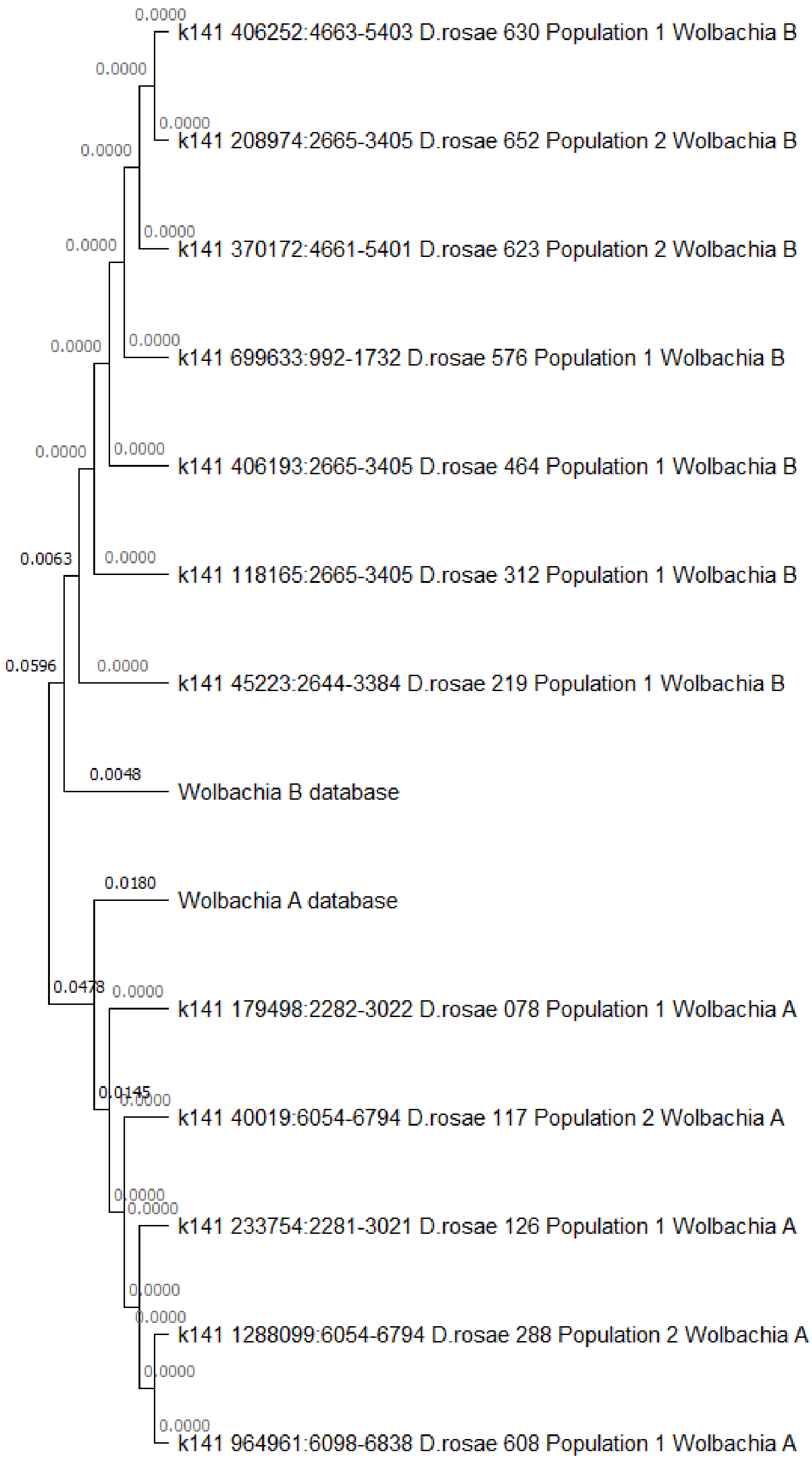

**Fig. S13**. Phyloogram of *Wolbachia* genotypes based on the *hcpA* gene (Wang et al. 2020) recovered from *Diplolepis rosae* Illumina sequences. Each sample corresponds to *Wolbachia* contig identifier: start–end position of the gene_corresponding *D.rosae* sample name_*D. rosae* population_*Wolbachia* supergroup. The phylogram was constructed using the Maximum likelihood statistical method (Tamura–Nei substitution model (default)) by MEGA 11 (Tamura et al. 2021).

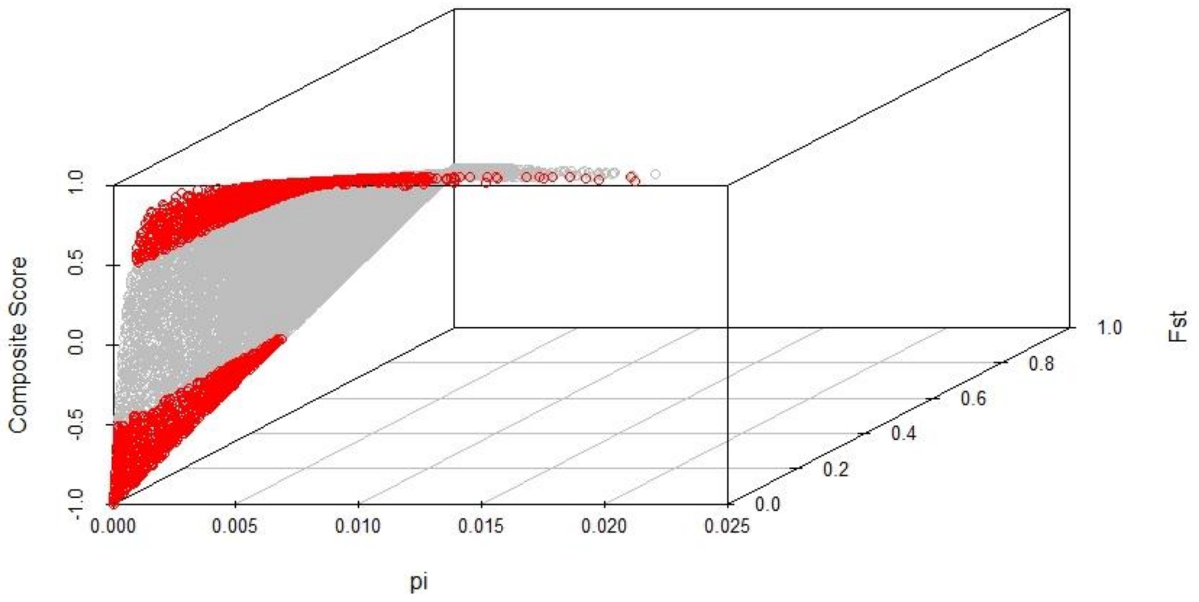

**Fig. S14**. Relationship between nucleotide diversity (pi), the fixation index Fst, and Composite Score (CS) equal to $(1 - Fst)*2(F(pi) - 0.5)$, where F(pi) is the cumulative distribution function. Red points represent CS outliers below −0.5 and above 0.5 corresponding to the genome regions of *Diplolepis rosae* examined in Gene Set Enrichment Analysis.

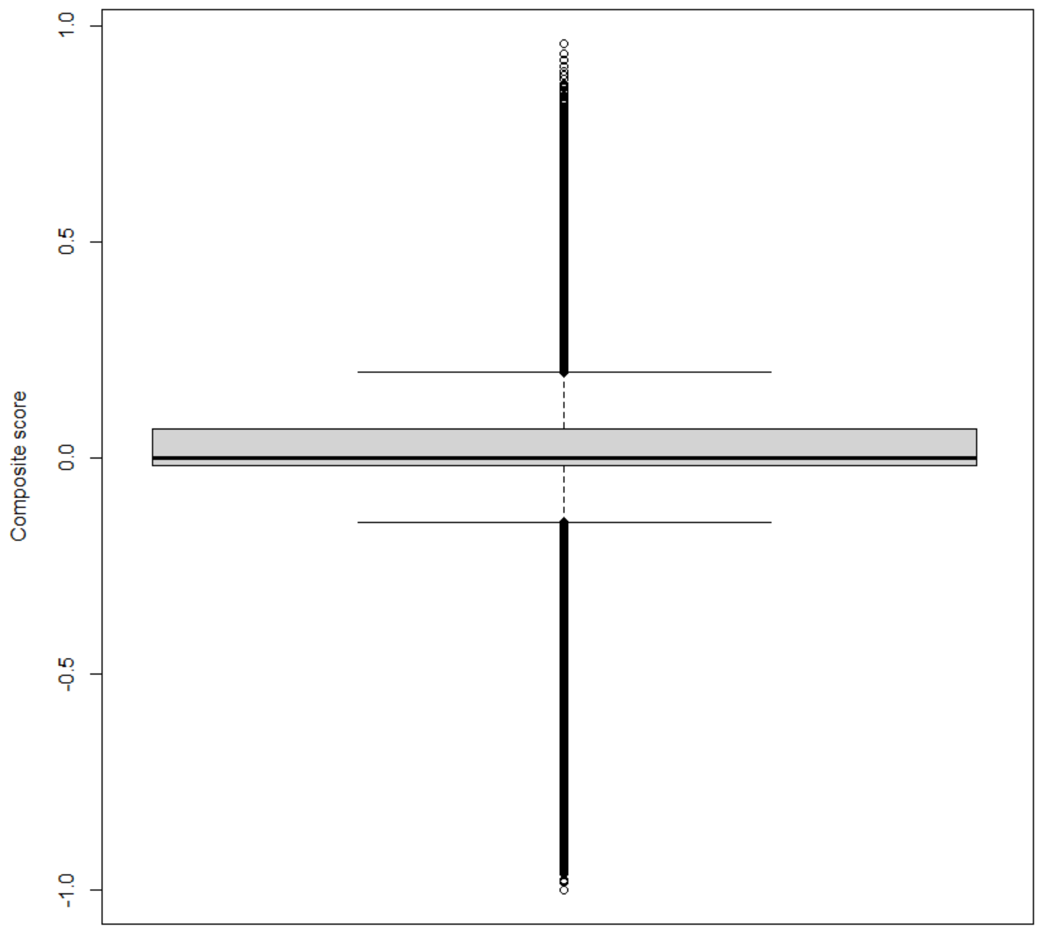

**Fig. S15**. Distribution of composite score (CS) values calculated for 10-kbp windows in the *Diplolepis rosae* genome. CS equals (1 − Fst)*2(F(π) − 0.5), where Fst is the fixation index calculated between the two *D. rosae* populations, π is the nucleotide diversity calculated for each *D. rosae* population, and F(π) is the cumulative distribution function.

### Protocols

**Protocol S1.** Polymerase chain reaction protocol using a *Diplolepis rosae* genetic marker.

1. For one sample in 200-μL plastic tubes prepare 25 μL of a solution containing:
    - 5 μL of 5X Taq Buffer with MgCl2 (Promega)
    - 0.5 μL of 10 mM dNTPs (Promega)
    - one unit of Taq DNA polymerase (0.2 μL) (Promega)
    - 0.7 μL of 10 mM forward primer †
    - 0.7 μL of 10 mM reverse primer †
    - 16.9 μL of nuclease-free water (Promega)
    - 1 μL of extracted DNA
2. PCR thermocycling profile:
    - 1 cycle of 2 min at 95 °C
    - 45 cycles of 45 sec at 95 °C, 45 sec at 65.5 °C, and 60 sec at 72 °C
    - final extension of 5 minutes at 72 °C
    - store at 4 °C until necessary

† *Information about the marker:*

Sequence:

GCATGAGATTGAGAAGCGGGAGTGGTGGCTTTGACGACTGCCACCCTCGAAGACTA
GAAGAAGGGCTTATTCCGCCGTCGTTTCAACCGGCGATGATCATTAAAAGTTTAAGC
GTCCCCTATTCAAGTACAGATTCAACGGCCGATTCAGATTCTACTATGGACGGCGCT
ATTTACTTTCTGCTCTCGTCAAACTTGCTAAGAGAAAAGAGAGAGAGCAATTATTGG
TAATTCGGTTTCATAAATTTCGGTAAAATGAATTTCAGGAAATATGGAATTTGGGGA
ATATTCATGTGCCCCTCCGAATAACTATGCGAAGTGTGTTCAGAAAGTAATGATAAA
ATCTTCAGCTGTATTCATTGTTTACTTTGTCCAGGCACGGCCAAGTCTTATAGGAGG
AATTGGCAAAATTTAGAACTTGGACGAAGGGATGAAGGGGGAAGTGCGGGGTGTTG
AGTAGTTATCTGGTCCCTGGTAGGTGGCTCTGGATATGATGACTAGATGTATCTGCG
GGGAAAGGGGATTTGAATAATGCAAACACGAGACAGTCTCAATTTTGGTTAGTTGC
TGCCAGTTATTTATGGAAAATGGATCGTAATAAATGCATTGTGCCAGCTTGTAATTC
ATTACGGAAGAATGAAAAAGTTCCAATACATCGACTTCCAAGAAATAGTGTCACTG
GACGTGAGTGGTTAGAGTTGTGCGGA

Primer design:

| OLIGO | start | len | tm | gc% | any_th | 3'_th | hairpin | seq |
|---|---|---|---|---|---|---|---|---|
| LEFT PRIMER | 7319 | 20 | 59.06 | 55.00 | 0.00 | 0.00 | 0.00 | |
| 5’ GCATGAGATTGAGAAGCGGG 3’ | | | | | | | | |
| RIGHT PRIMER | 8024 | 20 | 58.95 | 50.00 | 0.00 | 0.00 | 0.00 | |
| 5’ TCCGCACAACTCTAACCACT 3’ | | | | | | | | |

SEQUENCE SIZE: 8323

INCLUDED REGION SIZE: 8323

PRODUCT SIZE: 706, PAIR ANY_TH COMPL: 5.58, PAIR 3'_TH COMPL: 8.79

### Codes

**Code S1.** Scripts used to find the best scenario describing the demography of *Diplolepis rosae* by the continuous approximation approach.

```
python3
import numpy as np
import dadi

#General remarks
#Time T is given in units of 2*Neff generations
#Migration rate m12 is given in units of 2*Neff*mig12, mig12 is a
#fraction of individuals each generation in pop 1 that are new migrants #from
pop 2 rates. No migration m12=m21=0; simmetric migration m12=m21=m
#pts_0 = [20, 40, 60] is the number of grid points used in the calculation

#Importing data
dd = dadi.Misc.make_data_dict_vcf("varian_call_Drosae_ancestral_allele.vcf",
"pops.txt")
fs = dadi.Spectrum.from_data_dict(dd, ['pop1', 'pop2'], projections = [18, 16],
polarized=True)

#Joint polarized AFS
import pylab
pylab.figure(figsize=(10,10))
dadi.Plotting.plot_single_2d_sfs(fs, vmin = 1)
pylab.savefig('AFS', dpi=250)

#Single-population statistics
thetaW = fs.Watterson_theta()
pi = fs.pi()
D = fs.Tajima_D()

#Multi-population statistics
S = fs.S()
Fst = fs.Fst()

#split model: split into two populations of a specified size with migration
#nu1, nu2: population sizes after split
#T: time in the past of split
#m12, m21: migration

def split_mig(params, ns, pts):
      nu1,nu2,T,m12,m21 = params
      xx = dadi.Numerics.default_grid(pts
      phi = dadi.PhiManip.phi_1D(xx)
      phi = dadi.PhiManip.phi_1D_to_2D(xx, phi)
      phi = dadi.Integration.two_pops(phi, xx, T, nu1, nu2, m12=12, m21=m21)
      fs = dadi.Spectrum.from_phi(phi, ns, (xx,xx))
      return fs

#optimisation of parameters
my_extrap_func = dadi.Numerics.make_extrap_func(split_mig)
pts_0 = [20, 40, 60]
upper_bound = [2000, 2000, 0.5, 0.15, 0.15]
```

```
lower_bound = [500, 500, 0.1, 0.05, 0.05]
#set of parameters from which start optimisation
p0 = [1250, 1250, 0.25, 0.1, 0.1]
p0 = dadi.Misc.perturb_params(p0, fold=1, upper_bound=upper_bound, lower_bound=lower_bound)
print('Beginning optimisation')
popt = dadi.Inference.optimize(p0, fs, my_extrap_func, pts_0, lower_bound=lower_bound, upper_bound=upper_bound, verbose=len(p0))
print('Finished optimisation')
print('Best-fit parameters: {0}'.format(popt))

#Best-fit parameters after optimisation
#When assymetric migration:
#popt = [7.51053057e+02 1.98023428e+03 1.69649627e-01 5.28043606e-02 #1.48500000e-01]
#When symmetric migration m12=m21=m:
#popt = [1.02722156e+03 1.56986352e+03 2.17817079e-01 1.48500000e-01]
#When no migration:
#popt = [1.16747258e+03 1.99816894e+03 1.74841169e-01]

#isolation model: isolation with exponential pop growth
#s: proportion of individuals of pop 1 after split (pop 2 has size 1-s)
#nu1, nu2: final population sizes (after exponential growth)
#T: time in the past of split
#m12, m21: migration

def isol_mig(params, ns, pts):
    s,nu1,nu2,T,m12,m21 = params
    xx = dadi.Numerics.default_grid(pts)
    phi = dadi.PhiManip.phi_1D(xx)
    phi = dadi.PhiManip.phi_1D_to_2D(xx, phi)
    nu1_func = lambda t: s * (nu1/s)**(t/T)
    nu2_func = lambda t: (1-s) * (nu2/(1-s))**(t/T)
    phi = dadi.Integration.two_pops(phi, xx, T, nu1_func, nu2_func, m12=12, m21=21)
    fs = dadi.Spectrum.from_phi(phi, ns, (xx,xx))
    return fs

#optimisation of parameters
my_extrap_func = dadi.Numerics.make_extrap_func(isol_mig)
pts_0 = [20, 40, 60]
upper_bound = [0.9, 2000, 2000, 1.5, 1.5, 1.5]
lower_bound = [0.1, 500, 500, 0.05, 0.05, 0.05]
p0 = [0.5, 1000, 1000, 0.1, 1, 01]
p0 = dadi.Misc.perturb_params(p0, fold=1, upper_bound=upper_bound, lower_bound=lower_bound)
print('Beginning optimisation')
popt = dadi.Inference.optimize(p0, fs, my_extrap_func, pts_0, lower_bound=lower_bound, upper_bound=upper_bound, verbose=len(p0))
print('Finished optimisation')
print('Best-fit parameters: {0}'.format(popt))

#Best-fit parameters after optimisation
#When assymetric migration:
```

```
#popt = [8.21020680e-01 1.55906681e+03 1.20456650e+03 9.10803571e-01
#1.35621133e+00 7.18192319e-0]
#When symmetric migration m12=m21=m:
#popt = [1.00004257e-01 5.26620953e+02 8.98966318e+02 1.46967456e-01
#7.69940616e-01]
#When no migration:
#popt = [8.95917278e-01 6.99188738e+02 1.66546204e+03 4.53489214e-01]

#Isolation-with-migration model with exponential pop growth and a size change
prior to split
#nuPre: size after first size change
#TPre: time before split of first size change
#s: fraction of nuPre that goes to pop 1 => pop 2 has size nuPre*(1-s).)
#nu1, nu2: final population sizes
#T: Time in the past of split
#m12, m21: migration
#n1,n2: sample sizes of resulting spectrum

def isol_mig_pre(params, ns, pts):
      nuPre,TPre,s,nu1,nu2,T,m12,m21 = params
      xx = Numerics.default_grid(pts)
      phi = PhiManip.phi_1D(xx)
      phi = Integration.one_pop(phi, xx, TPre, nu=nuPre)
      phi = PhiManip.phi_1D_to_2D(xx, phi)
      nu1_0 = nuPre*s
      nu2_0 = nuPre*(1-s)
      nu1_func = lambda t: nu1_0 * (nu1/nu1_0)**(t/T)
      nu2_func = lambda t: nu2_0 * (nu2/nu2_0)**(t/T)
      phi = Integration.two_pops(phi, xx, T, nu1_func, nu2_func, m12=m12,
m21=m21)
      fs = Spectrum.from_phi(phi, ns, (xx,xx))
      return fs

#optimisation of parameters
my_extrap_func = dadi.Numerics.make_extrap_func(isol_mig_pre)
pts_0 = [20, 40, 60]
upper_bound = [2000, 6, 0.9, 1000, 1000, 1.5, 1.5, 1.5]
lower_bound = [500, 5, 0.1, 800, 800, 0.5, 0.5, 0.5]
p0 = [1000, 5.5, 0.5, 900, 900, 1, 1, 1]
p0 = dadi.Misc.perturb_params(p0, fold=1, upper_bound=upper_bound,
lower_bound=lower_bound)
print('Beginning optimisation')
popt = dadi.Inference.optimize(p0, fs, my_extrap_func, pts_0,
lower_bound=lower_bound, upper_bound=upper_bound, verbose=len(p0))
print('Finished optimisation')
print('Best-fit parameters: {0}'.format(popt))

#Best-fit parameters after optimisation
#When assymetric migration:
#popt = [7.64925622e+02 5.01183360e+00 7.53743684e-01 9.89999994e+02
#8.07999931e+02 8.54984055e-01 8.16222033e-01 9.00362495e-01]
#When symmetric migration m12=m21=m:
#popt = [9.88797106e+02 5.05000000e+00 8.73576179e-01 9.90000000e+02
#8.53851232e+02 7.73547199e-01 9.16075629e-01]
#When no migration:
```

```
#popt = [5.15031419e+02 5.05000000e+00 7.07834470e-01 9.90000000e+02 #8.08000000e+02 1.48500000e+00]

#bottlegrowth_split model: instantanous size change followed by exponential growth then split
#nuB: ratio of population size after instantanous change to ancient #population size
#nuF: ratio of contempoary to ancient population size
#T: time in the past at which instantaneous change happened and growth #began
#Ts: time in the past at which the two populations split

def bottlegrowth_split(params, ns, pts):
    nuB,nuF,T,Ts = params
    return bottlegrowth_split_mig((nuB,nuF,0,T,Ts), ns, pts)

#optimisation of parameters
my_extrap_func = dadi.Numerics.make_extrap_func(bottlegrowth_split)
pts_0 = [20, 40, 60]
upper_bound = [0.2, 3, 0.5, 0.5]
lower_bound = [0.05, 1, 0.05, 0.05]
p0 = [0.1, 2, 0.1, 0.1]
p0 = dadi.Misc.perturb_params(p0, fold=1, upper_bound=upper_bound, lower_bound=lower_bound)
print('Beginning optimisation')
popt = dadi.Inference.optimize(p0, fs, my_extrap_func, pts_0, lower_bound=lower_bound, upper_bound=upper_bound, verbose=len(p0))
print('Finished optimisation')
print('Best-fit parameters: {0}'.format(popt))

#Best-fit parameters after optimisation
#popt = [0.13041597 2.37120407 0.21729583 0.20394267]

#bottlegrowth_split_migration model: instantanous size change followed by exponential growth then split with migration
#nuB: ratio of population size after instantanous change to ancient #population size
#nuF: Ratio of contempoary to ancient population size
#m: migration
#T: time in the past at which instantaneous change happened and growth began
#Ts: Time in the past at which the two populations split

def bottlegrowth_split_mig(params, ns, pts):
      nuB,nuF,m,T,Ts = params
      xx = dadi.Numerics.default_grid(pts)
      phi = dadi.PhiManip.phi_1D(xx)
      if T >= Ts:
            nu_func = lambda t: nuB*numpy.exp(numpy.log(nuF/nuB) * t/T)
            phi = dadi.Integration.one_pop(phi, xx, T-Ts, nu_func)
            phi = dadi.PhiManip.phi_1D_to_2D(xx, phi)
            nu0 = nu_func(T-Ts)
            nu_func = lambda t: nu0*numpy.exp(numpy.log(nuF/nu0) * t/Ts)
            phi = dadi.Integration.two_pops(phi, xx, Ts, nu_func, nu_func, m12=m, m21=m)
```

```
    else:
        phi = dadi.PhiManip.phi_1D_to_2D(xx, phi)
        phi = dadi.Integration.two_pops(phi, xx, Ts-T, 1, 1, m12=m, m21=m)
        nu_func = lambda t: nuB*numpy.exp(numpy.log(nuF/nuB) * t/T)
        phi  =  dadi.Integration.two_pops(phi,  xx,  T,  nu_func,  nu_func,
m12=m, m21=m)

     fs = dadi.Spectrum.from_phi(phi, ns, (xx,xx))
     return fs

#optimisation of parameters
my_extrap_func = dadi.Numerics.make_extrap_func(bottlegrowth_split_mig)
pts_0 = [20, 40, 60]
upper_bound = [0.9, 5, 1, 5, 5]
lower_bound = [0.01, 0.1, 0.1, 0.1, 0.1]
p0 = [0.1, 1, 0.5, 1, 1]
p0    =    dadi.Misc.perturb_params(p0,    fold=1,    upper_bound=upper_bound,
lower_bound=lower_bound)
print('Beginning optimisation')
popt    =    dadi.Inference.optimize(p0,    fs,    my_extrap_func,    pts_1,
lower_bound=lower_bound, upper_bound=upper_bound, verbose=len(p0))
print('Finished optimisation')
print('Best-fit parameters: {0}'.format(popt))

#Best-fit parameters after optimisation
#popt = [0.05008461 1.93543999 0.18266133 0.12533342 0.81468721]

#likelihood
model = my_extrap_func(popt, ns, pts_0)
ll = dadi.Inference.ll_multinom(model, fs)
print(ll)

#likelihood ratio test
#my_extrap_func_1: extrapolation of more complex model to an infinitely #fine
grid
#ptc_1: number of grid points used for more complex model
#p1: optimised parameters for more complex model
#ll_0, ll_1: composite likelihood of a simplier and more complex model,
respectively
#bootstrapping for more complex model
chunks = dadi.Misc.fragment_data_dict(dd, chunk_size=15000)
boots = dadi.Misc.bootstraps_from_dd_chunks(chunks, Nboot=10, pop_ids=['pop0',
'pop1'], projections = [18, 16], mask_corners=True, polarized=True)
#Multiplicative adjustment to the likelihood ratio test statistic
adj  =  dadi.Godambe.LRT_adjust(my_extrap_func_1,  ptc_1,  boots,  p1,  fs,
nested_indices=[5], multinom = True, eps=0.01)
#adjusted D-statistics
D=adj*2*(ll_1 - ll_0)
print (D)
p=dadi.Godambe.sum_chi2_ppf(D, weights=(0, 1))
print(p)
```

**Code S2**. Scripts used to estimate the model parameters of *Diplolepis rosae* demography (approximate Bayesian computation approach).

***Simulations***

```
import numpy as np
import msprime
import tskit
import csv

header = ['Nanc', 'N1', 'N2', 'tbot', 'proportion', 'tsplit', 'mu', 'mean_pi1', 'var_pi1', 'mean_pi2', 'var_pi2', 'mean_Tajimas_D1', 'var_Tajimas_D1', 'mean_Tajimas_D2', 'var_Tajimas_D2', 'mean_d', 'var_d', 'mean_Fst', 'var_Fst']
f = open("simulations_Drosae.csv", 'w', encoding='UTF8', newline='')
writer = csv.writer(f, delimiter="\t")
writer.writerow(header)

#In a log uniform distribution, the log transformed random variable is #assumed to be uniformly distributed
#Thus logU(a, b) ~ exp(U(log(a), log(b))
#Thus, we could create a log-uniform distribution using numpy
#mu: mutation rate (per sequence length per generation)
#N1, N2: diploid size of ancestral and two populations of D. rosae, #respectively
#tbot: bottleneck time of ancestral population, in generations
#proportion: proportion of individuals after bottleneck compared to the #size of ancestral population
#tsplit: split time of the ancestral population to two populations of D. #rosae, in generations

i = 1
while i <= 1000000:
#Random parameters
    mu = np.random.uniform(1e-10, 9.99e-7)
    N1 = np.exp(np.random.uniform(5, 12, size=None))
    N2 = np.exp(np.random.uniform(5, 12, size=None))
    tbot = np.exp(np.random.uniform(5, 9, size=None))
    proportion = np.random.uniform(0, 1)
    tsplit = np.exp(np.random.uniform(5, 9, size=None))
    if tbot> tsplit:
#Demography
            demography = msprime.Demography()
            demography.add_population(name="Pop1", initial_size=N1)
            demography.add_population(name="Pop2", initial_size=N2)
            demography.add_population(name="Anc", initial_size=N1+N2)
            demography.add_population_split(time=tsplit, derived=["Pop1", "Pop2"], ancestral="Anc")
            demography.add_simple_bottleneck(time=tbot, proportion=proportion, population="Anc")
#Tree sequence
#Sequence length: D. rosae assembly length 497872038 bp => 10000 times shorter

            sequence_length = 49787
            ts = msprime.sim_ancestry(samples={"Pop1": 9, "Pop2": 8}, demography=demography, ploidy=2, random_seed=np.random.randint(1, 5000 + 1), sequence_length=sequence_length)
```

```
#Mutation model
            model=msprime.BinaryMutationModel()
#Mutations
            mutations = msprime.sim_mutations(ts, rate=mu, random_seed=np.random.randint(1, 5000 + 1), model=model)
#Statistics in each 1000 bp region
            num_windows = 50
            pi1 = mutations.diversity(sample_sets=mutations.samples(population=0), windows=np.linspace(0, sequence_length, num_windows +1))
            pi2 = mutations.diversity(sample_sets=mutations.samples(population=1), windows=np.linspace(0, sequence_length, num_windows +1))
            Tajimas_D1 = mutations.Tajimas_D(sample_sets=mutations.samples(population=0), windows=np.linspace(0, sequence_length, num_windows +1))
            Tajimas_D2 = mutations.Tajimas_D(sample_sets=mutations.samples(population=1), windows=np.linspace(0, sequence_length, num_windows +1))
            d = mutations.divergence(sample_sets=[mutations.samples(population=0), mutations.samples(population=1)], windows=np.linspace(0, sequence_length, num_windows +1))
            Fst = mutations.Fst(sample_sets=[mutations.samples(population=0), mutations.samples(population=1)], windows=np.linspace(0, sequence_length, num_windows +1))
            data = [N1, N2, tbot, proportion, tsplit, mu, np.mean(pi), np.var(pi), np.mean(pi1), np.var(pi1), np.mean(pi2), np.var(pi2), np.mean(Tajimas_D), np.var(Tajimas_D), np.mean(Tajimas_D1), np.var(Tajimas_D1), np.mean(Tajimas_D2), np.var(Tajimas_D2), np.mean(d), np.var(d), np.mean(Fst), np.var(Fst)]
            writer.writerow(data)
            i += 1
      else:
            pass
```

#### *Approximate Bayesian Computation (abc package in R)*

A) Calculation of summary statistics (observed data, obs.txt): see **Code S8**

B) Parameter file (params.txt): simulation of N1, N2, mu, proportion, tbot, tsplit

C) Sim_sumstas file (sim_sumstats.txt): simulations of pi, Tajima's D, dxy, and Fst

```
#Import the data
obs <- read.table("obs.txt", header=TRUE)
params <- read.table("params.txt", header=TRUE)
sim_sumstats <- read.table("sim_sumstats.txt", header=TRUE)
#Goodness-of-fit
fit <- gfit(target=obs, sumstat=sim_sumstats, nb.replicate=100)
#Cross-validation
```

```
cross <- cv4abc(param=params, sumstat=sim_sumstats, abc.out = NULL, nval=100, tols=c(0.005, 0.01, 0.05), method="neuralnet")

#Parameter inference

rej <- abc(target=obs, param=params, sumstat=sim_sumstats, tol=0.01, method="neuralnet")

#Diagnostic plot

plot <- gfitpca(target=obs, sumstat=sim_sumstats, index="D", cprob=0.05)
```
